# Structural conformations of intrinsically disordered proteins of podocyte slit-diaphragm

**DOI:** 10.1101/2025.03.04.641475

**Authors:** Hari Krishnan Padmanaban, Sneha Bheemireddy, Sandeep KN Mulukala, Nageswara Rao Dunna, Anil Kumar Pasupulati

## Abstract

The slit-diaphragm (SD), a specialized junction between the foot processes of neighboring podocytes, regulates the permselectivity of glomerular filtration. It functions as a size- and charge-selective molecular sieve, ensuing protein-free urine. SD comprises several proteins such as nephrin, NEPH1, podocin, CD2AP, and TRPC6. Nephrin and NEPH1 are extracellular proteins that bridge the gap between foot processes, whereas podocin and CD2AP are adapter proteins. The intrinsically disordered regions (IDRs) of SD proteins play a crucial role in assembling these proteins as a macromolecular complex. Mutations in these proteins disrupt SD integrity, leading to a nephrotic syndrome characterized by heavy proteinuria. The structural details of each protein of this macromolecular complex are poorly detailed. We employed molecular docking and molecular dynamics simulations to investigate the structural dynamics of SD proteins. Our findings reveal that SD proteins exhibit partner-specific conformational adaptations driven by short linear motifs and molecular recognition features. CD2AP interacts transiently with nephrin but forms a more stable complex with podocin. NEPH1 and nephrin interact via extracellular immunoglobulin domains while maintaining dynamic intracellular contacts. Podocin preferentially interacts with nephrin over NEPH1, and distinct subunit-sharing mechanisms emerge, where CD2AP and TRPC6 may simultaneously associate with a single podocin subunit. Mutational analysis reveals that disease-associated variants, including CD2AP (P532S), podocin (R138Q), and nephrin (G1161V), enhance local stability but restrict the flexibility of IDR and impair SD assembly. Our study provides evidence of dynamic subunit sharing within the SD complex, offering new insights into the assembly of SD in health and disease.

## Introduction

At the heart of renal filtration lies the tri-layered glomerular filtration barrier (GFB), composed of inner endothelium, the middle glomerular basement membrane, and the outer layer of podocytes (Anil Kumar et al., 2014). Podocytes are unique cells with a large body and complex foot processes (FP). The FP that arose from primary and secondary processes of podocytes interdigitate with FP of neighboring podocytes by forming a modified tight junction known as slit-diaphragm (SD), a size, shape, and charge-selective barrier at the blood-urine interface. SD is formed by intricate network of proteins that include podocin, nephrin, CD2-associated protein (CD2AP), nephrin-like 1 (NEPH1), podocalyxin, transient receptor potential cation channel (TRPC6), among others (Fukasawa et al., 2009; Mulukala Narasimha et al., 2019; Mulukala et al., 2016; Pätäri-Sampo et al., 2006). Negatively charged surface anionic proteins within the foot processes contribute to the electrostatic barrier function of the SD, effectively restricting the passage of negatively charged macromolecules, such as serum albumin and immunoglobulins, into the glomerular filtrate. Together with other members of the GFB, SD of podocytes prevent filtration of proteins and large molecules and allow only water, electrolytes, and metabolic waste products (urea, uric acid, and creatinine) into primary urine, thus maintaining the body’s fluid levels and electrolyte balance (Anil Kumar et al., 2014; Mukhi et al., 2017).

NEPH1 and nephrin, both transmembrane proteins, form critical homophilic and heterophilic extracellular contacts between adjacent podocyte foot processes, while podocin and CD2AP are cytoplasmic and serve as adapter proteins and contribute to the structural integrity of the SD (Grahammer et al., 2016; Reiser et al., 2005). Biophysical studies have shown podocin interacts with CD2AP and nephrin, while TRPC6 associates with nephrin and podocin, forming a highly interconnected network (Schwarz et al., 2001; Shih et al., 2001). Evidence suggests that SD proteins may assemble into higher-order oligomeric structures (Qadri et al., 2024; Watanabe et al., 2000), but the molecular mechanisms driving these interactions remain unclear. Computational models indicate that intrinsically disordered regions (IDRs) mediate key interfacial interactions among SD proteins. However, how these IDRs influence binding specificity, conformational adaptability, and the impact of disease-causing mutations remains poorly understood.

Monogenic mutations in podocyte proteins are strongly linked to nephrotic syndrome (NS), a renal disease marked by severe proteinuria, hypoalbuminemia, and edema (Benoit et al., 2010; Sadowski et al., 2015). NS is classified into steroid-sensitive (SSNS) and steroid-resistant (SRNS) forms, with SRNS often progressing to end-stage renal disease (ESKD). Mutations in podocin (*NPHS2*) frequently cause SRNS, accounting for ∼30% of cases, while CD2AP mutations are associated with focal segmental glomerulosclerosis (FSGS) (Gigante et al., 2009; Sadowski et al., 2015). *NEPH1* and nephrin (*NPHS1*) mutations disrupt transepithelial permeability, with nephrin variants leading to congenital nephrotic syndrome (CNS) of the Finnish type, whereas *TRPC6* mutations dysregulate calcium signaling, exacerbating podocyte damage (Gigante et al., 2011; Kestilä et al., 1998; Solanki et al., 2019). The genetic heterogeneity of SRNS underscores the urgent need for mechanistic insights into SD protein interactions to inform targeted therapeutic strategies (Preston et al., 2019; Smith et al., 2007). Notably, early structure prediction algorithms have struggled to accurately capture the structural-functional relationships of SRNS-associated mutations (∼40% accuracy in CASP12).

Earlier studies establish interactions among selected SD proteins, including podocin, CD2AP, NEPH1, TRPC6, and nephrin (Benzing, 2004; Grahammer et al., 2016; Kocylowski et al., 2022; Reiser et al., 2005; Schwarz et al., 2001). While significant progress has been made in identifying individual protein-protein interactions, the precise molecular mechanisms governing their assembly into a supramolecular complex remain largely unclear. An important unresolved question is how SD proteins coordinate their interactions to establish stability for SD. Equally critical, yet poorly understood, is the impact of disease-associated mutations, particularly those linked to nephrotic syndromes such as SRNS, on these protein complexes’ assembly and structural integrity. These mutations frequently occur within IDRs or at key interfacial residues, which pose significant analytical challenges due to their dynamic nature and the limitations of earlier structure prediction models and experimental techniques. Consequently, there is a pressing need to unravel the interplay between wild-type (WT) and mutant forms of SD proteins to comprehensively understand how structural perturbations disrupt the fine assembly of SD that eventually manifest in proteinuria. This study aims to address these knowledge gaps by leveraging advancements in protein structure prediction and molecular dynamics (MD) simulations. Using AlphaFold-predicted models (Jumper et al., 2021) for podocin, CD2AP, NEPH1, TRPC6, and nephrin, we systematically generated all plausible heterodimeric complexes of these proteins through molecular docking. This approach allowed us to explore the interaction landscape and identify potential binding interfaces critical for complex formation. To further dissect the molecular mechanisms of these interactions, we subjected the docked complexes to extensive MD simulations, examining the proteins’ WT and mutant forms.

## Methods

### Molecular docking studies

We obtained the full-length structures of podocin, CD2AP, NEPH1, TRPC6, and nephrin from the AlphaFold database (https://alphafold.ebi.ac.uk/). Since the complete structures of these proteins remain unresolved, we truncated the models to retain only their interacting regions for molecular docking and molecular dynamics (MD) simulations. We retrieved protein sequences from UniProt (www.uniprot.org) and assessed their secondary structure content using the Phyre2 web server. For molecular docking, we used HADDOCK 2.4 (https://wenmr.science.uu.nl/haddock2.4/) and ClusPro 2.0 (https://cluspro.bu.edu/home.php) (Honorato et al., 2024; Kozakov et al., 2017). We docked CD2AP-nephrin and Nephrin-NEPH1 cytoplasmic complexes using HADDOCK in an information-driven manner, while we employed ClusPro for blind docking of the remaining complexes (Barletta et al., 2003; Shih et al., 2001). We selected the highest-ranked docking models for further analysis.

### MD simulations

We conducted MD simulations using GROMACS (ver. 2022.6) (Abraham et al., 2015) with the CHARMM force field (Brooks et al., 2009). We placed each protein complex in a 1 nm^3^ cubic box, solvated it in a TIP3 water system, and neutralized it with 0.15M NaCl using CHARMM-GUI (Jo et al., 2008). We minimized the systems using 2000 steps, followed by stepwise heating from 0 to 310 K, equilibration with backbone restraints, and production runs without restraints at 1 atm pressure. We set the short-range non-bonded interaction cutoff to 12 Å, with a shifting function at 10 Å. We calculated long-range electrostatic interactions using the Particle-Mesh Ewald method (Darden et al., 1993). We simulated WT protein-protein complexes for 100 ns and extracted interacting interfaces for further analysis using PAE (predicted aligned error) and pLDDT plots.

### Mutation studies

To identify disease-associated mutations, we queried dbSNP (Sherry et al., 2001), gnomAD (Karczewski et al., 2020), ClinVar (Landrum et al., 2014), COSMIC (Tate et al., 2019), and Ensembl (Martin et al., 2023). We assessed mutation pathogenicity using fathmm (Functional Analysis through Hidden Markov Models) (https://fathmm.biocompute.org.uk/index.html), LIST-S2 (https://list-s2.msl.ubc.ca/), PANTHER (https://www.pantherdb.org/), and Mutation Assessor (Malhis et al., 2020; Shihab et al., 2013; Tang & Thomas, 2016). We classified a mutation as pathogenic if it met the following criteria: predicted as damaging by FATHMM, scored above 0.85 in LIST-S2, ranked as medium or high risk in Mutation Assessor, and marked as probably damaging by PANTHER. We cross-referenced pathogenicity scores to pinpoint SRNS-associated mutations and mapped them onto 100 ns simulated WT complexes. We introduced these mutations into protein-protein complexes and performed an additional 50 ns MD simulations. A comprehensive list of all complexes can be found in **Table S1**.

### Visualization and analysis

We visualized simulation snapshots and videos using VMD and ChimeraX (Humphrey et al., 1996; Meng et al., 2023). We analyzed non-covalent interactions with PyContact, applying an H-bond distance cutoff of 2.5 Å and an angle cutoff of 120° (Scheurer et al., 2018). We defined protein-protein interactions based on a mean interaction score above one. We characterized MD trajectories using GROMACS functions, extracting insights into secondary structure changes, mean movement, and stable complexation (Van Der Spoel et al., 2005). We calculated binding free energies using the MM-GBSA (molecular mechanics with generalized Born and surface area solvation) method (Kumari et al., 2014).

## Results and Discussion

### CD2AP-Nephrin interaction dynamics implicate a transient nature

Previous investigations using co-immunoprecipitation and localization studies have shown that the C-terminus of CD2AP interacts with the cytoplasmic tail of nephrin. However, the specific residues involved in these interactions remain unidentified. We conducted MD simulations to explore the interaction between CD2AP and the cytoplasmic domain of nephrin (modeled structures included in this study are illustrated in **Figure 1a- f**). We evaluated the predicted models using PAE (predicted aligned error) and pLDDT plots **(Figure S1 and Table 1)**, and the detailed workflow is outlined in **Figure 2**. The CD2AP-nephrin heterodimer maintained contact for approximately 60% of the simulation time (∼65 ns of direct interactions and ∼35 ns of transient interactions, **Movie S1**). CD2AP’s coiled-coil domain (residues 569–639) **(Figure 3a)** interacted with a short α- helix in nephrin (residues 1213–1220). Additionally, two short linear motifs (SLiMs) in CD2AP (residues 527–536 and 544–556) engaged with corresponding SLiMs in nephrin (residues 1201–1212 and 1221–1239) **(Table S2)**, contributing to interaction stability.

**Figure 1:**
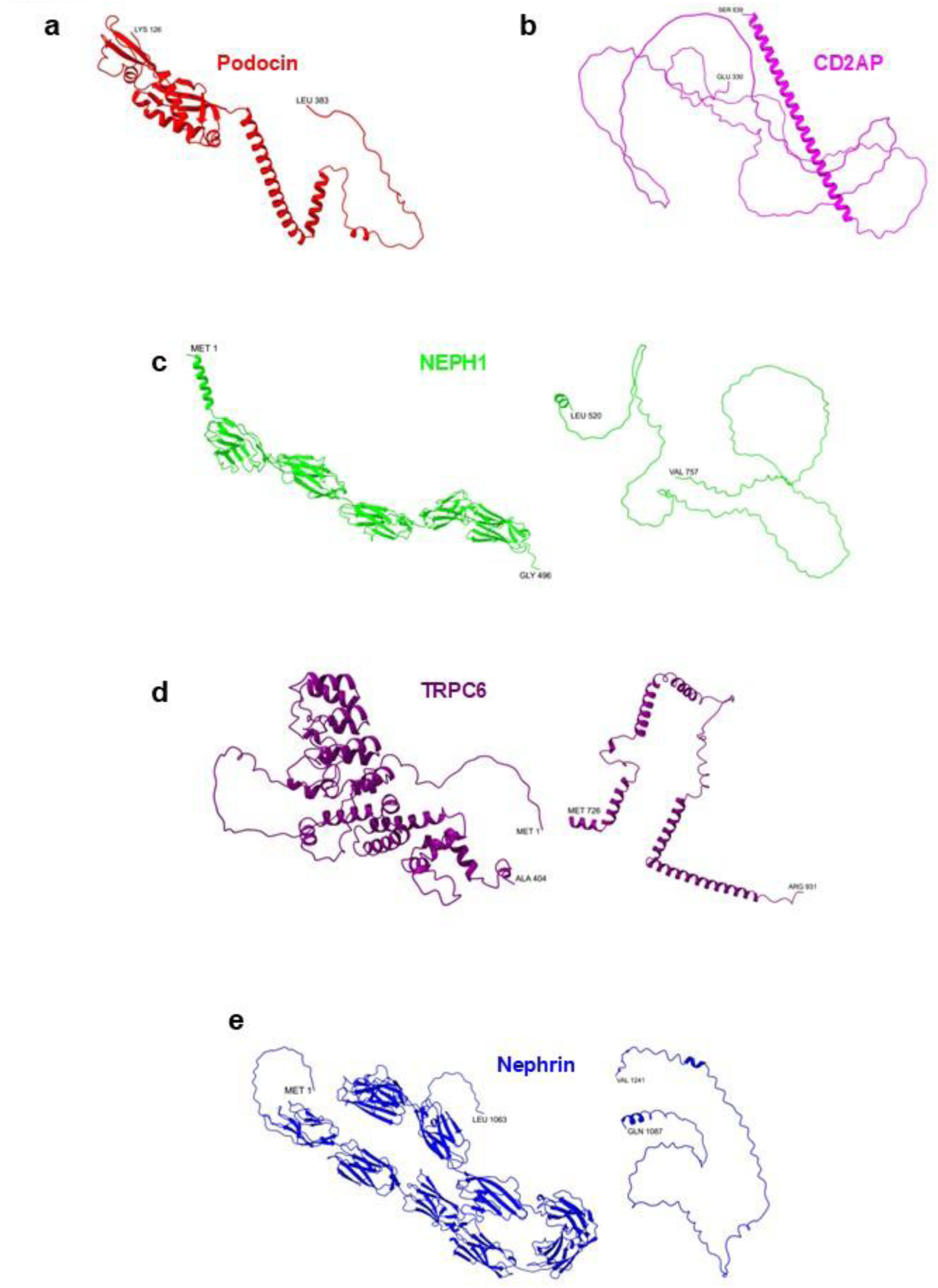
AlphaFold-predicted models of slit-diaphragm (SD) proteins included in the study. **a.** Podocin: Residues 126-383, which includes the PHB domain, are shown, excluding the N-terminal and transmembrane regions. **b.** CD2AP: Residues 330-639, incorporating the C-terminal coiled-coil domain, are displayed. **c.** NEPH1: The extracellular domain (1- 496) containing five Ig-like domains (1-5) and disordered cytoplasmic domain (520-757) are presented, excluding the transmembrane region. **d.** TRPC6: N-terminal containing domain (1-404) incorporating four ANK repeats (1-4), a TRP domain (5), and C-terminal containing domain (726-931) are considered, omitting the 6-pass transmembrane region. **e.** Nephrin: The extracellular domain (1-1063) containing nine Ig-like domains (1-9), an FN3 domain (10), and disordered cytoplasmic domain (1087-1241) are displayed, excluding the transmembrane region. **f.** HSP27: The full-length protein model (residues 1-205) is presented. All proteins are color-coded and represented in a cartoon format.

**Figure 2:**
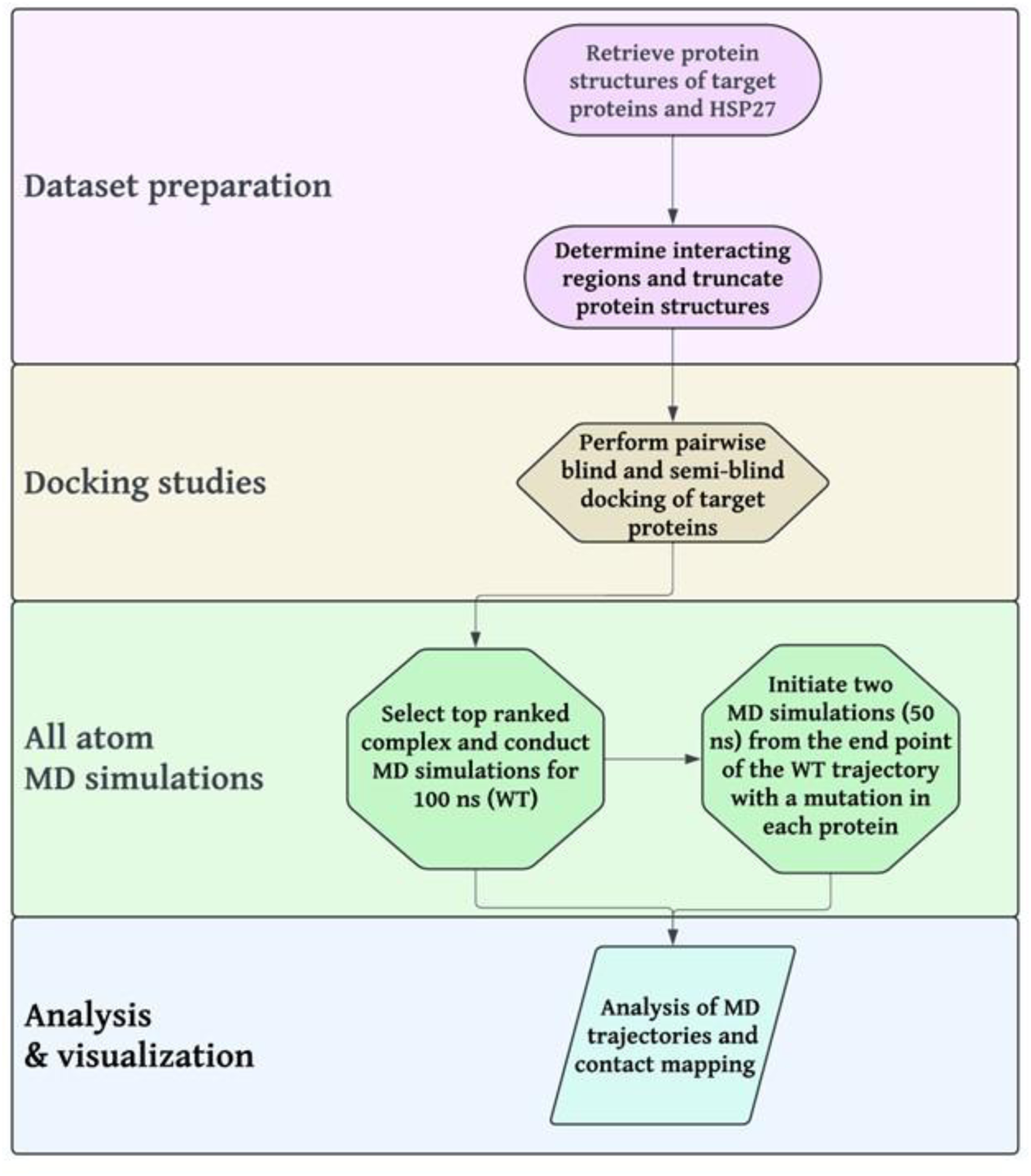
Schematic overview of methodology. The workflow begins with the extraction and truncation (as required) of AlphaFold-predicted models of the target proteins. These processed models are then subjected to blind or semi-blind docking to generate all plausible heterodimer interactions. The resulting complexes undergo all-atom MD simulations and extensive analysis to assess their stability and interactions.

**Figure 3:**
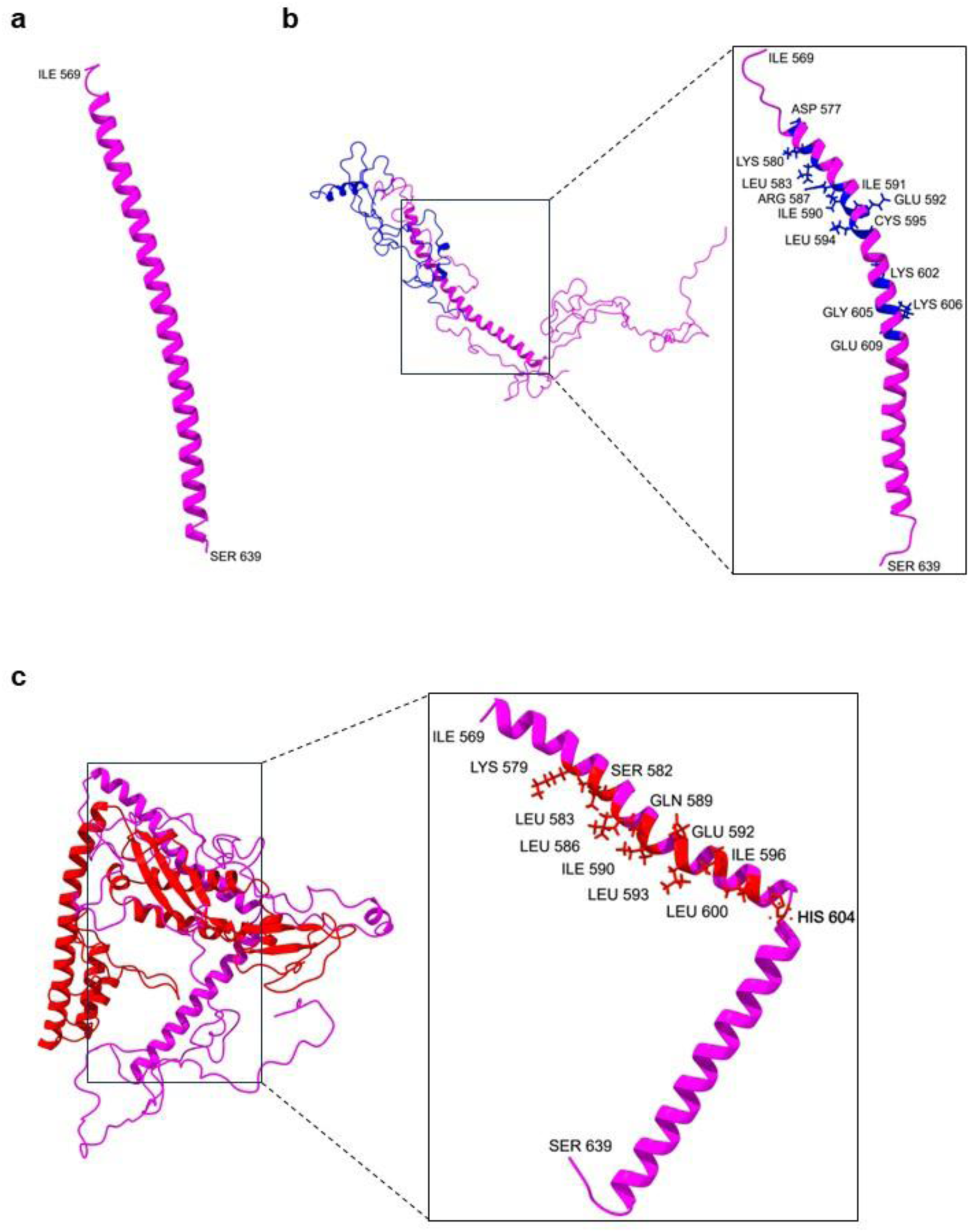
Partner-specific folding of CD2AP’s coiled-coil domain. **a**. Initial structure of CD2AP’s C-terminal coiled-coil domain (residues 569–639) before MD simulations. CD2AP is displayed as a magenta cartoon. **b.** CD2AP coiled-coil domain after 100 ns of MD simulations with nephrin (blue cartoon), showing an RMSD of 4.5 Å over 66 Cα atoms. The residues in the CD2AP coiled-coil domain that interact with nephrin are highlighted in blue licorice representations. **c.** CD2AP coiled-coil domain after 100 ns of MD simulations with podocin (red cartoon), showing a significantly higher RMSD of 11.8 Å over 69 Cα atoms. Red licorices in CD2AP’s coiled-coil domain actively interact with podocin.

**Table 1:** Physical characteristics of selected SD proteins.

| Gene / Chromosome | Protein | Uniprot ID | KDa | AA | % Disorder | % Disallowed residues (Rama. plot) | Selected point-mutations |
| --- | --- | --- | --- | --- | --- | --- | --- |
| *NPHS2* / 1(q25.2) | Podocin | Q9NP85 | 42 | 383 | 47 | 3.3 | *p.R138Q*<br>*p.R238S*<br>*p.R322Q* |
| *CD2AP* / 6(p12.3) | CD2AP | Q9Y5K6 | 70 | 639 | 59 | 12.1 | *p.P532S* |
| *KIRREL1* / 1(q23.1) | NEPH1 | Q96J84 | 83 | 757 | 38 | 6.1 | *p.R440C*<br>*p.S573L* |
| *TRPC6* / 11(q22.2) | TRPC6 | Q9Y210 | 102 | 931 | 13 | 2.2 | *p.R175W*<br>*p.R895C* |
| *NPHS1* / 19(q13.12) | Nephrin | O60500 | 136 | 1241 | 24 | 1.3 | *p.C265R*<br>*p.G1161V* |

The CD2AP coiled-coil domain showed a root mean square deviation (RMSD) of 4.5 Å over 66 Cα atoms **(Figure 3b)**, indicating moderate flexibility in the interaction. While RMSD values below 2–3 Å typically suggest rigid, stable complexes, the observed deviation indicates that the CD2AP-nephrin interface retains some conformational adaptability, characteristic of interactions involving IDRs. Binding energy analysis (ΔG = −67 kcal/mol; **Figure S2a**) places this interaction within moderate-to-strong protein-protein interactions, comparable to other SD protein interactions such as podocin-CD2AP. However, despite the favorable binding energy, the interaction remained transient, with CD2AP disengaging intermittently throughout the simulation. These observations suggest that CD2AP-nephrin binding may serve a regulatory or signaling role rather than forming a stable structural complex, consistent with the dynamic nature of IDR-mediated interactions. The transient nature of this interaction may be critical for maintaining SD plasticity and reorganization, particularly in response to podocyte signaling cues or mechanical stress.

### Podocin induces conformational changes in CD2AP

We next analyzed the association between CD2AP and podocin. Previous studies have shown that podocin interacts with CD2AP through its C-terminal region (Schwarz et al., 2001), yet the specific binding region in CD2AP remains unclear. Our simulations provided critical information about exact CD2AP regions involved in this interaction. In the CD2AP-podocin complex, CD2AP’s interacting region was primarily composed of IDRs (residues 332–568), whereas podocin’s interaction was confined to its structured prohibitin homology (PHB) domain (residues 131–286) **(Table S3)**. Over the 100 ns simulation, CD2AP’s coiled-coil domain underwent structural rearrangement upon engaging with podocin, with residues 579–604 forming stable contacts that induced a pronounced bend at residues 603–606 **(Figure 3c, Movie S2)**. This conformational change directed residues 607–638 inward toward the complex center, leading to an RMSD of 11.8 Å over 69 aligned Cα atoms. Additionally, three SLiMs in CD2AP (residues 448–461, 467–480, and 500–513) engaged podocin, further stabilizing the complex. Podocin’s PHB domain (residues 131–282) exhibited minor conformational adjustments, reinforcing interaction stability **(Figure 4b)**.

**Figure 4:**
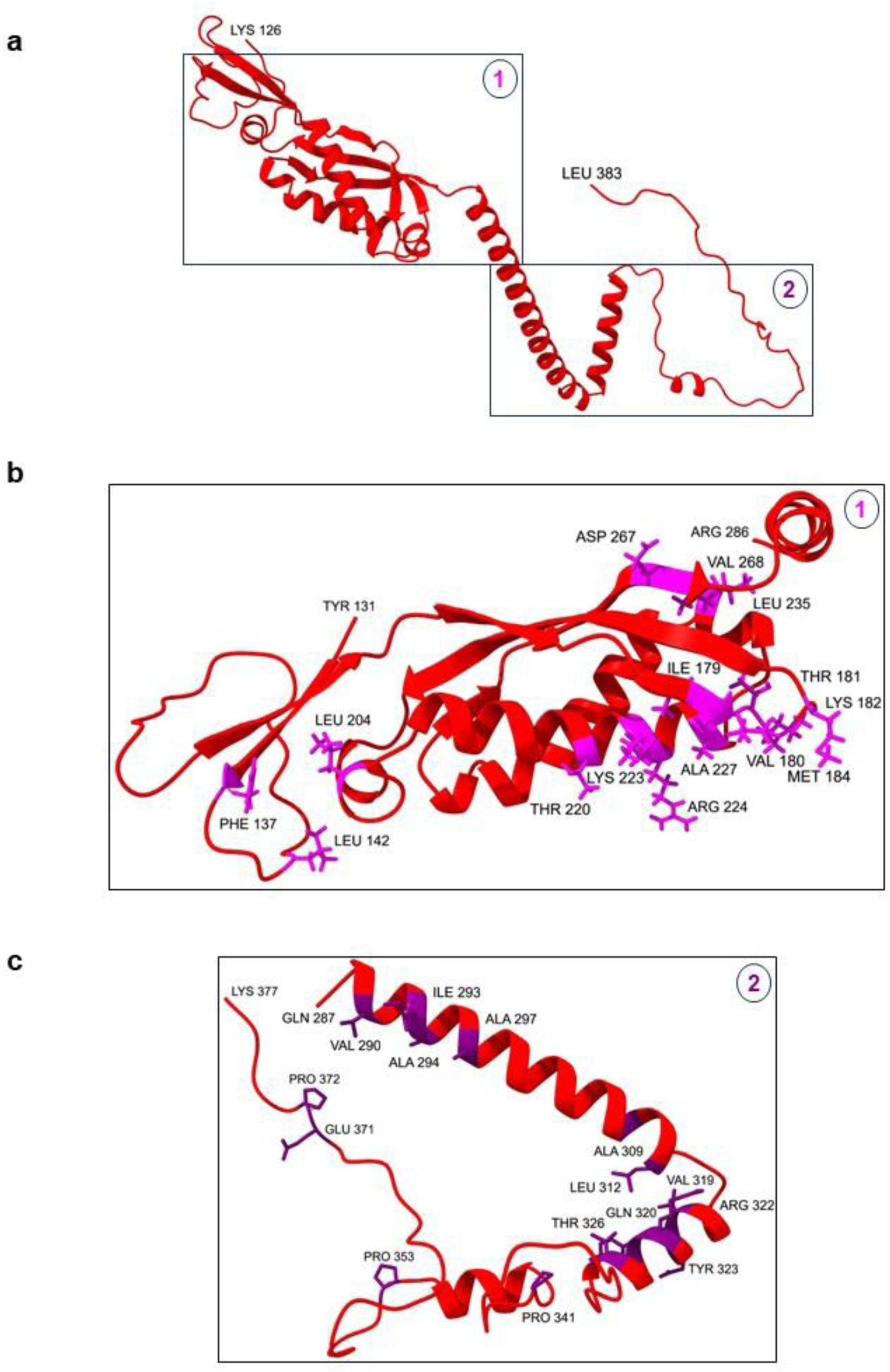
Podocin interacts with CD2AP and TRPC6 via distinct subregions of a single subunit. **a**. Initial structure of podocin’s C-terminal region (residues 126–383), encompassing the PHB domain, before MD simulations. Podocin is shown as a red cartoon for clarity, while water molecules and ions are omitted from the visualization. **b.** Podocin model (residues 131–286) after 100 ns of MD simulations with CD2AP, highlighting the interacting residues as magenta licorices. **c.** The podocin model (residues 287–377) after 100 ns of MD simulations with TRPC6. The residues of podocin interacting with TRPC6 are shown as purple licorice representations.

A ΔG of −103 kcal/mol was calculated for the CD2AP-podocin complex **(Figure S2b)**, suggesting a stronger interaction between CD2AP and podocin than between CD2AP and nephrin (ΔG = −67 kcal/mol). Interestingly, CD2AP residues 579–604 are predicted to participate in interactions with both podocin and nephrin, indicating that CD2AP may preferentially bind podocin based on observed binding energies and structural changes. This overlap suggests CD2AP may engage with each protein via different subunits. While interactions in CD2AP’s disordered regions overlapped between nephrin and podocin, the inherent flexibility of IDRs and the lower confidence in structural predictions necessitate caution in interpreting these interactions as stable or functionally significant.

### Mutations alter the binding affinity and structural dynamics of CD2AP-nephrin and CD2AP-podocin complexes

Following the initial 100 ns WT simulations, we introduced one mutation per protein per complex to assess their impact on CD2AP-nephrin and CD2AP-podocin interactions (see Methods for mutation-selection details). Introducing the P532S mutation in CD2AP increased contact points in both complexes **(Table S2)** and restricted the movement of CD2AP’s IDRs, which are dynamic in the WT CD2AP-podocin complex. In the CD2AP- nephrin complex, the G1161V mutation in nephrin significantly hindered the movement of CD2AP’s coiled-coil domain and nephrin’s C-terminal residues. Visual analysis showed that G1161V induced an α-helix-to-coil transition in nephrin’s 1087–1098 region **(Movie S1)**, suggesting that this mutation alters nephrin’s structural integrity and flexibility. The R138Q mutation in podocin enhanced contact points by 57% in the CD2AP-podocin complex, further stabilizing the interaction.

Interestingly, the P532S mutation in CD2AP and the R138Q mutation in podocin both facilitated the involvement of prolines at positions 420–422 of CD2AP, which are part of an Src homology 3 (SH3) binding motif. This is particularly significant because SH3 motifs have high affinity for SH3 domains, which mediate protein-protein interactions and intracellular signaling pathways. The P532S mutation in CD2AP restricted the flexibility of IDRs, while R138Q in podocin enhanced contact formation by 57%, increasing the stability of the CD2AP-podocin complex. This suggests that P532S strengthens CD2AP’s interaction network, while R138Q enhances podocin’s binding affinity, potentially altering SD protein complex formation.

Our data indicate that these mutations effectively doubled the binding affinities of the CD2AP-nephrin and CD2AP-podocin complexes, leading to stronger but more rigid interactions. While this increased stability might reinforce SD assembly, it could also limit the dynamic adaptability necessary for SD function. The reduced flexibility of SD proteins may impair the ability of podocyte foot processes to respond to glomerular hydrostatic forces, potentially contributing to disease-associated dysfunction.

For further insights into the structural effects of these mutations, including changes in root mean square deviation (RMSD), radius of gyration (Rg), and root mean square fluctuation (RMSF) please refer to supplementary figures **(Figures S3 and S4)**.

### The structural integrity of extracellular nephrin-NEPH1 complex

As mentioned in the introduction, nephrin and NEPH1 bridge neighboring podocyte foot processes via their extracellular domains, forming the core structural support of the SD. To examine the stability and interaction dynamics of this complex, we performed MD simulations of the extracellular portions of nephrin and NEPH1. The complex remained structurally stable over 100 ns, with an RMSD of 8 Å, calculated over 893 aligned Cα atoms **(Figure 5a)**. The interaction interface primarily consisted of immunoglobulin (Ig) domains from both nephrin and NEPH1, with nephrin residues 143–234, 740–832, and 838–939 and NEPH1 residues 21–115 and 120–216 being critical for binding **(Table S4)**.

**Figure 5:**
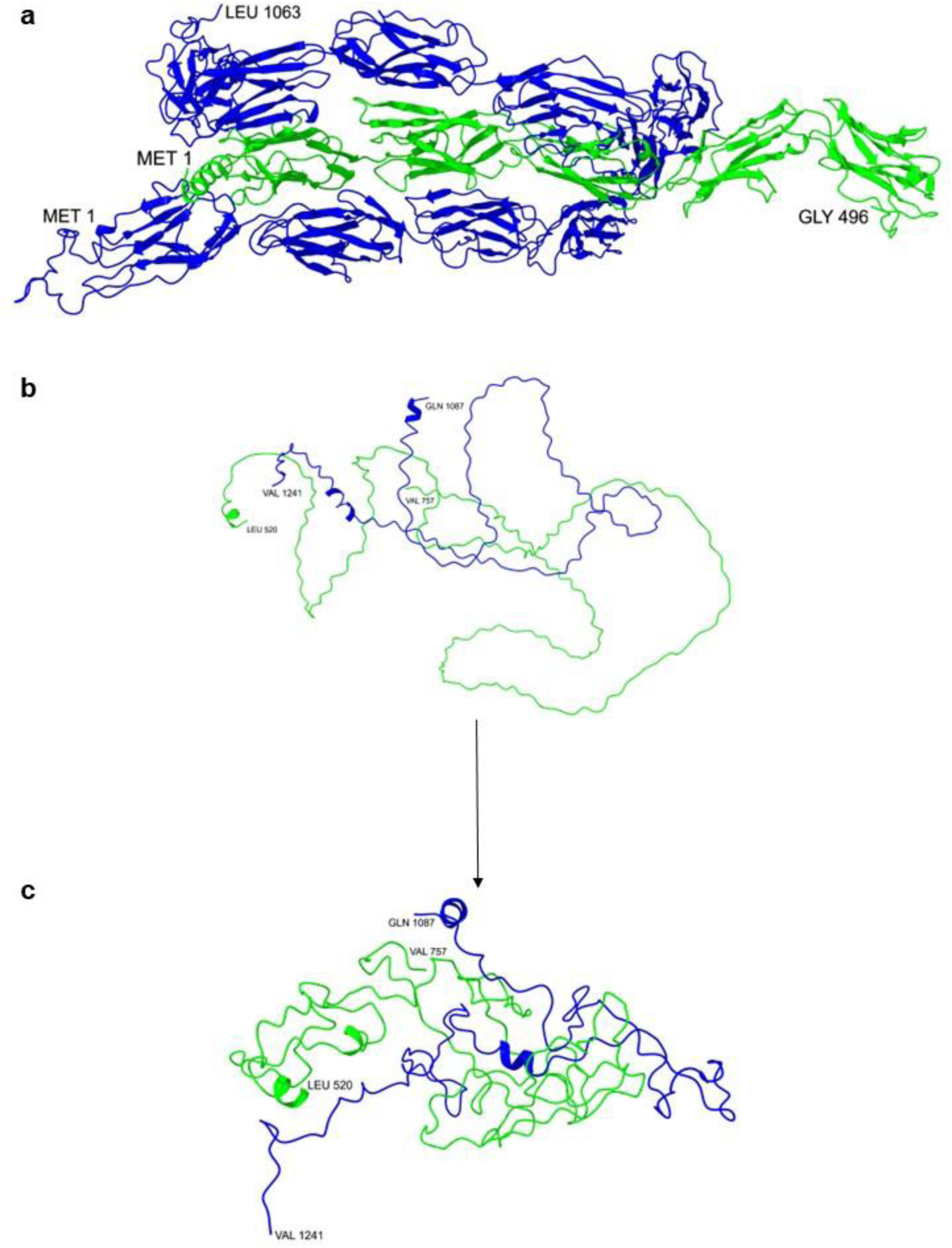
Cytoplasmic flexibility of the Nephrin-NEPH1 complex contributes to the rigidity of its extracellular contact. **a**. Model of the extracellular nephrin-NEPH1 complex after 100 ns of MD simulations, demonstrating a stable conformation. **b.** The initial model of the cytoplasmic Nephrin-NEPH1 complex shows both proteins in their native, extended conformations before MD simulations. **c.** Model of the cytoplasmic Nephrin-NEPH1 complex after 100 ns of MD simulations, showing a more compact conformation. NEPH1 and nephrin are represented as green and blue cartoons, respectively, to distinguish their contributions to the complex. Water molecules and ions are excluded from the visualization for clarity.

Interestingly, the Ig domains of nephrin contained residues associated with congenital nephrotic syndrome (CNS) of the Finnish type, suggesting a potential structural basis for disease-linked mutations in this region. However, the SRNS-associated residue 440 in NEPH1 did not contribute directly to the binding interface, indicating that its pathogenic effect may arise from disruptions in intracellular signaling rather than extracellular complex stability. Additionally, the N-terminal α-helix of NEPH1 (residues 11–20) played a key role in interacting with nephrin, reinforcing the importance of secondary structural elements in SD protein assembly **(Table S4)**. Apart from nephrin’s fibronectin type 3 (FN3) domain (residues 943–1038) and a flexible disordered region (residues 1039– 1063), the nephrin-NEPH1 complex remained stable throughout the simulation **(Movie S3)**. These findings highlight the critical role of the extracellular nephrin-NEPH1 interaction in maintaining SD integrity and function.

### Cytoplasmic interaction dynamics of Nephrin-NEPH1 complex

We extended our analysis to understand the intracellular interactions between NEPH1 and nephrin. Following docking, both proteins were primarily disordered and extended **(Figure 5b)**, but after 100 ns of simulation, the complex became more compact, with a decreased Rg **(Figure 5c, Movie S4)**. The predicted ΔG for the extracellular nephrin-NEPH1 complex was −62 kcal/mol **(Figure S2c)**, while the intracellular complex exhibited a stronger binding affinity with a ΔG of −82 kcal/mol **(Figure S2d)**. The intracellular complex showed a significant RMSD of 40 Å over 393 Cα atoms, reflecting the dynamic nature of the disordered regions. Further, we identified three SLiMs (residues 1107-1119, 1124-1136, and 1228-1235) and an MoRF (residues 1167-1218) at the interaction interface of nephrin-NEPH1 **(Table S5)**. While the last 41 residues of nephrin are predicted to interact with CD2AP, some residues in this region were found to engage NEPH1. Notably, despite being part of the immunoglobulin superfamily, the absence of significant sequence similarity in the cytoplasmic domains of NEPH1 and nephrin was striking.

### Impact of mutations on cytoplasmic and extracellular nephrin-NEPH1 complexes

Mutational analysis revealed that monogenic mutations in the cytoplasmic domains (NEPH1-S573L and nephrin-G1161V) tripled the binding affinities compared to the WT complex. In contrast, mutations in the extracellular domains (nephrin-C265R and NEPH1- R440C) resulted in only modest improvements (48% and 50% increases in interactions, respectively) **(Table S4)**. Specifically, S573L and G1161V increased contacts by 38% and 39% in the cytoplasmic complex **(Table S5)**. These mutations also reduced flexibility in specific regions, notably residues 650-750 in NEPH1 and 1120-1160 in nephrin, indicating a more stable interaction in the cytoplasmic domain. In contrast, extracellular interactions remained relatively stable, highlighting their robustness. Overall, the simulations distinguished the flexible intracellular region from the stable extracellular domain, revealing how mutations in the disordered cytoplasmic regions impair movement, similar to observations in the CD2AP complexes. Further details on the trajectory analysis of these complexes can be found in **Figures S5 and S6**. As discussed earlier, adapter proteins interact with the cytoplasmic domains of NEPH1 and nephrin. To further explore these interactions, we analyzed how the cytoplasmic domains of NEPH1 and nephrin interact with podocin, a major adapter protein in podocytes.

### Preferential binding of podocin with nephrin over NEPH1

Next, we analyzed the interactions between the cytoplasmic regions of podocin-nephrin and podocin-NEPH1 complexes. Podocin primarily interacted with NEPH1 and nephrin via its PHB and helical (271-332) domains **(Figure 6a)**, while NEPH1 (ΔG of −80 kcal/mol) and nephrin (ΔG of −95 kcal/mol) remained largely disordered **(Movies S5 and S6)**.

**Figure 6:**
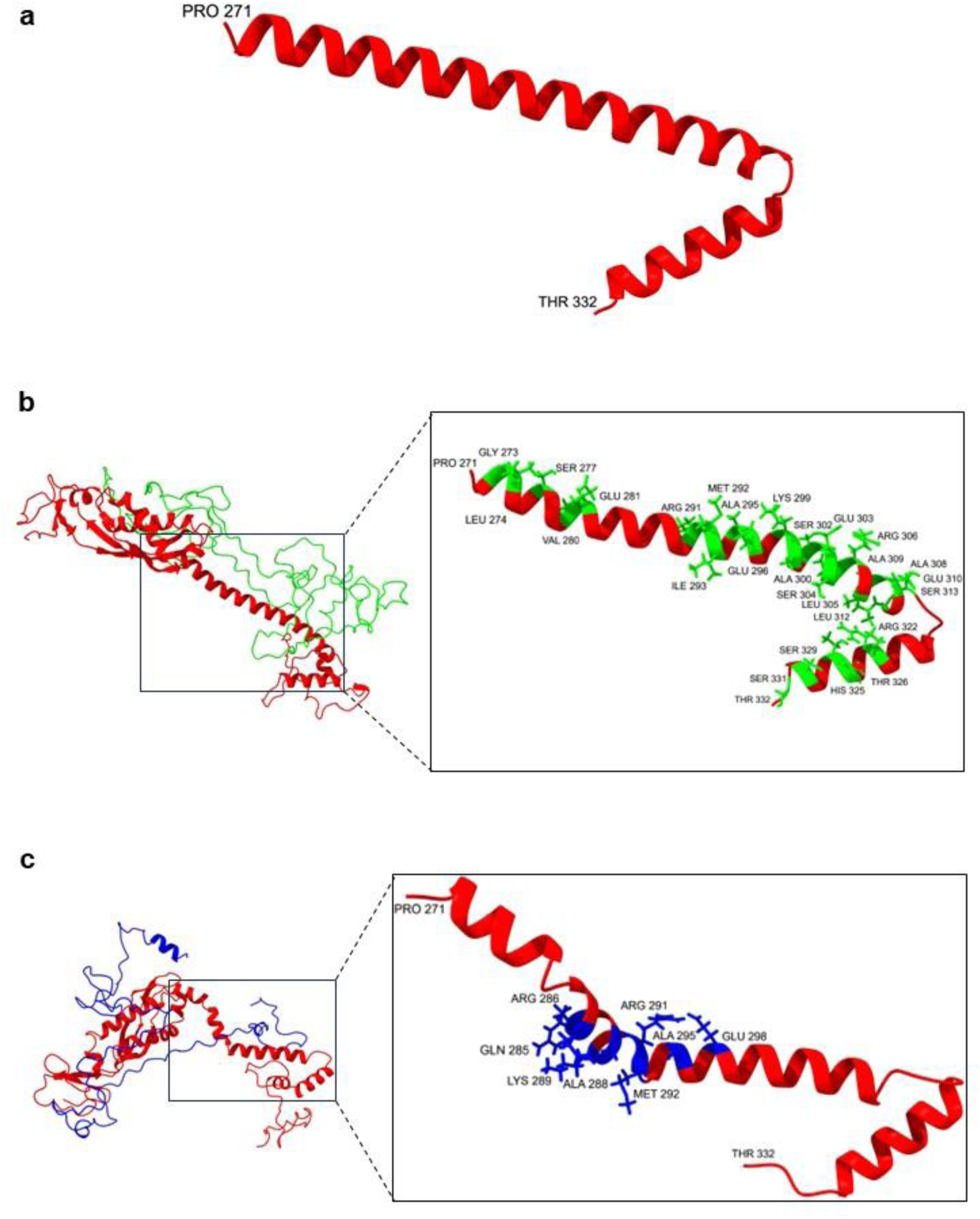
Nephrin induces more significant conformational changes in podocin compared to NEPH1. **a**. Initial model of podocin’s C-terminal helical region (residues 271-332) before MD simulations. Podocin is depicted as a red cartoon. **b.** After 100 ns of MD simulations with NEPH1 (green cartoon), the podocin’s C-terminal helical region model shows an RMSD of 1.6 Å across 55 Cα atoms, indicating minimal conformational change. **c.** After 100 ns of MD simulations with nephrin (blue cartoon), the podocin’s C-terminal helical region model shows a larger RMSD of 6.6 Å across 62 Cα atoms, reflecting significant conformational alteration. The residues of podocin, interacting with NEPH1 and nephrin, are highlighted in green and blue licorice representations, respectively.

After 100 ns of simulation with NEPH1, podocin’s helical domain exhibited minimal conformational change, with an RMSD of 1.6 Å over 55 aligned Cα atoms **(Figure 6b)**, indicating stable structural integrity. In contrast, NEPH1’s IDR changed from an extended to a compact state. We identified several SLiMs (542-551, 576-581, 586-594, 710-716, 736-742, 747-757) and a MoRF (609-636) in NEPH1 that were crucial for its interaction with podocin **(Table S6)**. NEPH1’s MoRF (609-636) and SLiM (736-742) interacted with podocin, similar to its interaction with nephrin. Notably, V180 residue in podocin, which also interacts with CD2AP, was also involved in binding with NEPH1 **(Table S6)**.

Interaction with nephrin induced a distinct bend in podocin’s helical domain (residues 285- 298), with an RMSD of 6.6 Å over 62 aligned Cα atoms **(Figure 6c)**, slightly altering podocin’s conformation as compared to its conformation in the podocin-NEPH1 complex. It is possible that IDRs 269-272, 312-316, 328-344 & 351-383 in podocin provide adaptive flexibility for podocin to interact in distinct confirmations with nephrin and NEPH1. Our simulations revealed that only eight residues of podocin’s helical domain interacted with nephrin, instead of twenty-eight residues that interacted with NEPH1 **(Figures 6b and 6c)**. Notably, in a similar pattern to the cytoplasmic NEPH1 region in the NEPH1-podocin complex, the nephrin’s disordered regions adopted a compact conformation upon binding to podocin **(Figure S8c, Movie S6)**. Interestingly, nephrin’s interfaces with NEPH1 and podocin did not overlap, suggesting that podocin and NEPH1 may share a nephrin subunit, while CD2AP might interact with a separate subunit. Our results indicate that the podocin-NEPH1 complex exhibited a slightly stronger binding affinity than the podocin-nephrin complex **(Figure S2f)**, highlighting the differential affinities of podocin for nephrin and NEPH1.

### Mutations in podocin, NEPH1, and nephrin alter conformational dynamics

We introduced three individual mutations—R238S in podocin, S573L in NEPH1, and G1161V in nephrin—to assess their impact on the stability and interactions of the podocin-NEPH1 and podocin-nephrin complexes. Each mutation was introduced while keeping its partner in the WT state. The R238S (podocin) and S573L (NEPH1) mutations significantly improved the energetic stability of the podocin-NEPH1 complex, enhancing binding affinity and increasing contact formation by 17% and 22%, respectively, compared to the WT podocin-NEPH1 complex **(Table S6, Figure S2e)**. The R238S mutation in podocin significantly restricted NEPH1’s movement, while the S573L mutation in NEPH1 had a similar effect on podocin **(Figures S7e and S7f)**. In the podocin-nephrin complex, the R238S and G1161V mutations increased contact formation by 30% and 42%, respectively, compared to the WT podocin-nephrin complex **(Table S7, Figure S2f)**. Interestingly, the R238S mutation provided greater flexibility to nephrin’s residues 1176- 1184, while G1161V increased flexibility in nephrin’s residues 1110-1117 and 1127-1138 **(Figure S8f)**. The introduction of R238S, S573L, and G1161V led to more compact conformations in the cytoplasmic regions of NEPH1 and nephrin when interacting with podocin **(Movies S5 and S6)**. In the podocin-NEPH1 complex, mutations reduced the flexibility of IDRs, whereas, in the podocin-nephrin complex, some SLiMs in nephrin gained more flexibility after introducing mutations. Refer **Figures S7 and S8** for detailed trajectory analysis.

### Potential role of podocin and nephrin in TRPC6 channel gating

We next examined the interaction between podocin, nephrin, and TRPC6. Previous studies showed that TRPC6 interacts with podocin via its C-terminal and with nephrin via its N-terminal (Reiser et al., 2005). Our analysis of the podocin-TRPC6 interaction revealed that podocin’s interface primarily consists of IDRs **(Figure 4c)**, while TRPC6 contains helical domains. Two SLiMs (333-344 & 366-377) in podocin **(Table S8)** induced conformational changes in TRPC6. TRPC6, initially in an extended conformation, became more compact upon interaction with podocin, as shown by changes in the radius of gyration **(Figure S9c)** and visualized in **Movie S7**. IDRs in TRPC6 (755-770, 789-796, 814-840, 849-854, 875-885) enabled these conformational alterations. Three specific helices (732-744, 769-788 & 795-812) in TRPC6 were observed in its interacting interface with podocin. A helical stretch (287-326), a MoRF (333-356), and an SLiM (366-377) of podocin interacted with TRPC6. Notably, podocin residues 287-377 exclusively interact with TRPC6, while residues 131-286 are reserved for CD2AP binding, with no overlap between the interfaces suggesting that CD2AP and TRPC6 can simultaneously share a podocin subunit.

The interaction between podocin and TRPC6 exhibited a binding affinity of −110 kcal/mol, while the TRPC6-nephrin interaction was stronger at −135 kcal/mol **(Figures S2g and S2h)**. Despite the cytoplasmic nephrin being completely disordered, its interaction with TRPC6 was more energetically favorable than podocin’s, highlighting the critical role of IDRs in nephrin in facilitating its interaction with TRPC6. TRPC6 helped nephrin to adopt a more compact structure **(Movie S8)**. TRPC6’s interface with nephrin included two SLiMs (13-20, 209-216), a MoRF (40-88), two ankyrin (ANK) repeats (163-189, 218-247), a TRP domain (253-315), and a helix (331-345) while, three SLiMs (1116-1129, 1136-1142 & 1155-1162) and a MoRF (1174-1211) of nephrin were a part of this interface **(Table S9)**. Nephrin’s residues 1201-1211 interacted with TRPC6 and CD2AP, though the nephrin-TRPC6 complex was more stable. Interestingly, the nephrin-CD2AP interaction was energetically costly, requiring twice the energy of the nephrin-TRPC6 interaction.

### Influence of mutations on TRPC6, podocin, and nephrin interactions

We introduced mutations in the TRPC6, podocin, and nephrin complexes to assess their impact on binding and dynamics. R895C in TRPC6’s C-terminal, which interacts with podocin, and R175W in TRPC6’s N-terminal, which interacts with nephrin, both enhanced the binding energy of their respective complexes **(Figures S2g and S2h)**. R322Q (podocin) improved the podocin-TRPC6 interactions, while R895C had no such effect **(Table S8)**. R322Q also increased the flexibility of podocin and TRPC6 (residues 768- 810), while R895C restricted their movement. In the TRPC6-nephrin complex, R175W was introduced in TRPC6, and G1161V was introduced in nephrin. These mutations increased interactions by 13% and 20%, respectively **(Table S9)**. G1161V (nephrin) reduced flexibility in both proteins, while R175W (TRPC6) enhanced the movement of nephrin’s residues 1091-1109. Detailed trajectory analysis is provided in **Figures S9 and S10**.

### Summary & Future Perspectives

Our study provides a comprehensive structural analysis of interactions of key proteins of SD, emphasizing the critical role of overlapping SLiMs and MoRFs in stabilizing the assembly of the complex. We identified distinct subunit-sharing mechanisms, where CD2AP and TRPC6 likely interact with the same podocin subunit, while NEPH1 and nephrin engage separate podocin subunits. These observations suggest that podocin should exist as an oligomer to maintain interactions with its partner proteins in the SD complex. Notably, our earlier study revealed that podocin exists as a higher-order oligomer (Mulukala et al., 2020). Podocin’s interfaces with NEPH1 and nephrin significantly overlapped, whereas its interactions with CD2AP and TRPC6 were distinct and non-overlapping. CD2AP’s interfaces with podocin and nephrin also exhibited some overlap, supporting the idea that CD2AP-podocin interactions are more direct, whereas CD2AP-nephrin interactions remain transient. These insights refine our understanding of SD protein organization, mainly how structural adaptability and selective interactions contribute to podocyte function.

Moreover, our findings reveal that mutations within IDRs can significantly alter binding affinity, disrupt native interactions, and reduce the flexibility required for SD protein function. The conformational changes in podocin and CD2AP upon binding different partners suggest that protein dynamics play a crucial role in SD assembly and maintenance. These results have direct implications for understanding the molecular basis of nephrotic syndrome. Future studies should expand on these findings by investigating potential homo-oligomeric states and larger biological assemblies, as SD proteins may function within higher-order complexes in a cellular environment. Additionally, while our simulations provided valuable insights, incorporating multiple initial conformations and iterative simulations will further enhance the robustness and accuracy of structural predictions. Building on this work, future experimental studies such as x-ray crystallography and cryo-EM could validate these computational findings and refine our understanding of SD protein interactions in disease contexts. Nevertheless, the major hurdle to pursuing experimental studies is the challenges of expressing and purifying SD proteins.

## Supporting information

Table S2 - CD2AP - Nephrin complex

Table S3 - CD2AP - Podocin complex

Table S4 - N.Nephrin - N.NEPH1 complex

Table S5 - C.NEPH1 - C.Nephrin complex

Table S6 - Podocin - NEPH1 complex

Table S7 - Podocin - Nephrin complex

Table S8 - Podocin - TRPC6 complex

Table S9 - TRPC6 - Nephrin complex

## Acknowledgements

AKP acknowledges the University of Hyderabad for the IoE grant and ICMR-DHR grant to visit Harvard Medical Center, Boston, USA.

