## Supplemental Data (Tables & Figures) for "Structural conformations of intrinsically disordered proteins of podocyte slit-diaphragm"

**Table S1: Cumulative summary of protein-protein simulations: trajectories and durations.** At a physiological pH of 7, all systems were protonated, solvated with explicit TIP3 water molecules, and supplemented with 0.15 M NaCl.

| S.No. | WT complex | Time of trajectory | Modified complex | Time of trajectory |
|---|---|---|---|---|
| 1 | CD2AP - Nephrin complex | 0 – 100 ns | CD2AP (*P532S*) – Nephrin | 101 – 150 ns |
| | | | CD2AP – Nephrin (*G1161V*) | |
| 2 | CD2AP - Podocin complex | 0 – 100 ns | CD2AP (*P532S*) – Podocin | 101 – 150 ns |
| | | | CD2AP – Podocin (*R138Q*) | |
| 3 | N.Nephrin – N.NEPH1 complex | 0 – 100 ns | Nephrin (*C265R*) – NEPH1 | 101 – 150 ns |
| | | | Nephrin – NEPH1 (*R440C*) | |
| 4 | C.NEPH1 – C.Nephrin complex | 0 – 100 ns | NEPH1 (*S573L*) – Nephrin | 101 – 150 ns |
| | | | NEPH1 – Nephrin (*G1161V*) | |
| 5 | Podocin – NEPH1 complex | 0 – 100 ns | Podocin (*R238S*) – NEPH1 | 101 – 150 ns |
| | | | Podocin – NEPH1 (*S573L*) | |
| 6 | Podocin – Nephrin complex | 0 – 100 ns | Podocin (*R238S*) – Nephrin | 101 – 150 ns |
| | | | Podocin – Nephrin (*G1161V*) | |
| 7 | Podocin – TRPC6 complex | 0 – 100 ns | Podocin (*R322Q*) – TRPC6 | 101 – 150 ns |
| | | | Podocin – TRPC6 (*R895C*) | |
| 8 | TRPC6 – Nephrin complex | 0 – 100 ns | TRPC6 (*R175W*) – Nephrin | 101 – 150 ns |
| | | | TRPC6 – Nephrin (*G1161V*) | |

#### Supplementary figure 1

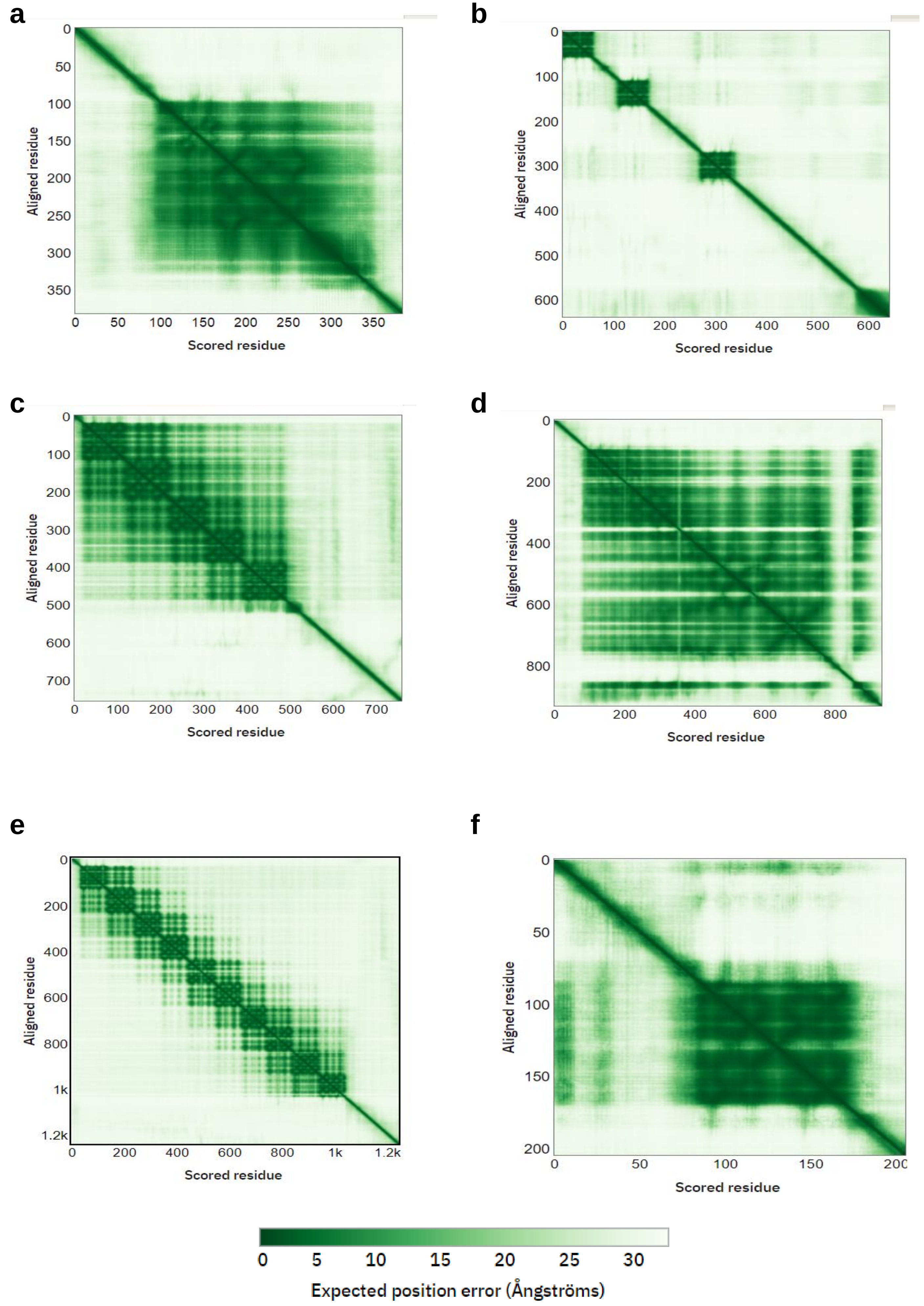

#### Supplementary figure 2

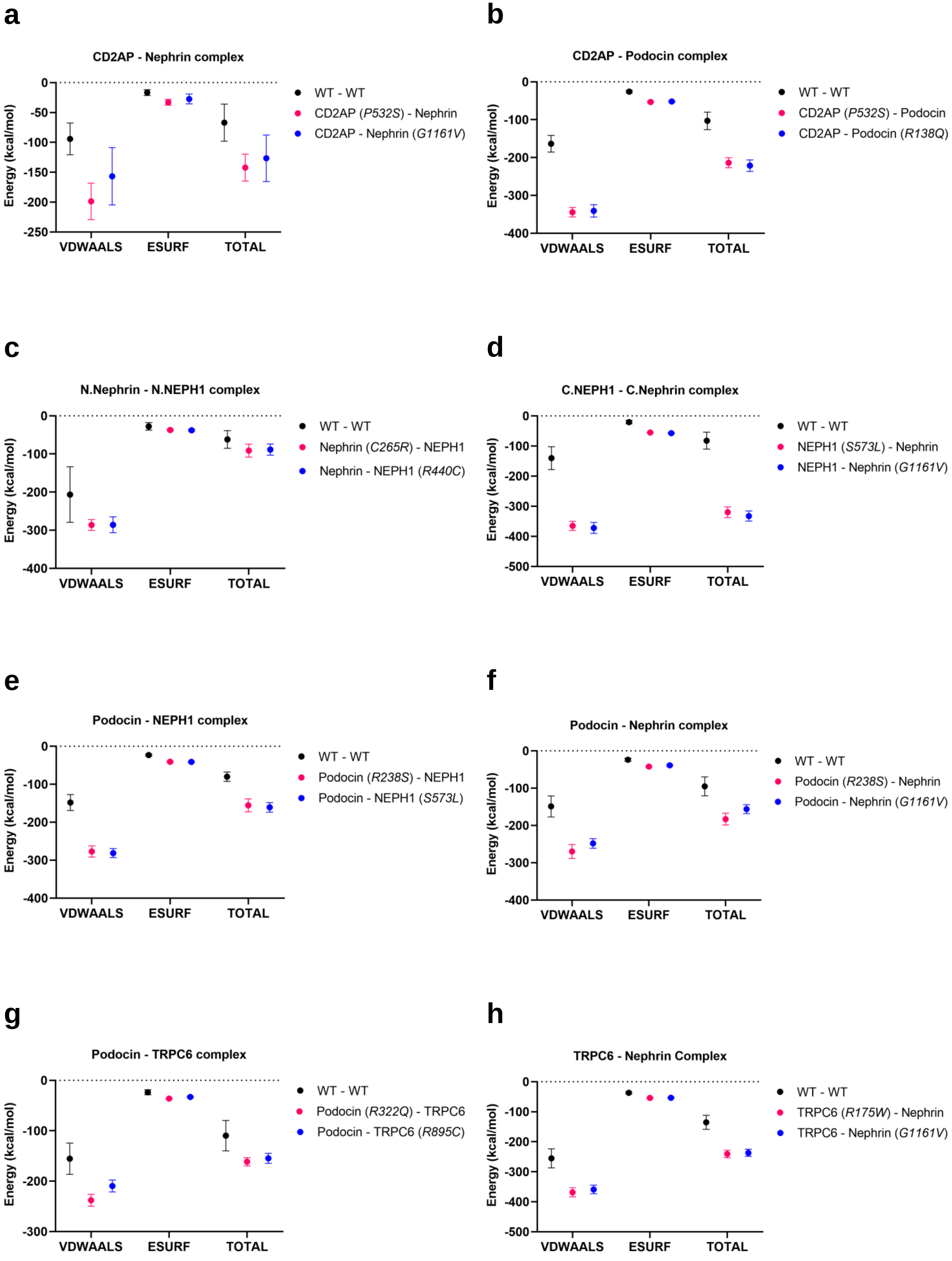

### Supplementary figure 3

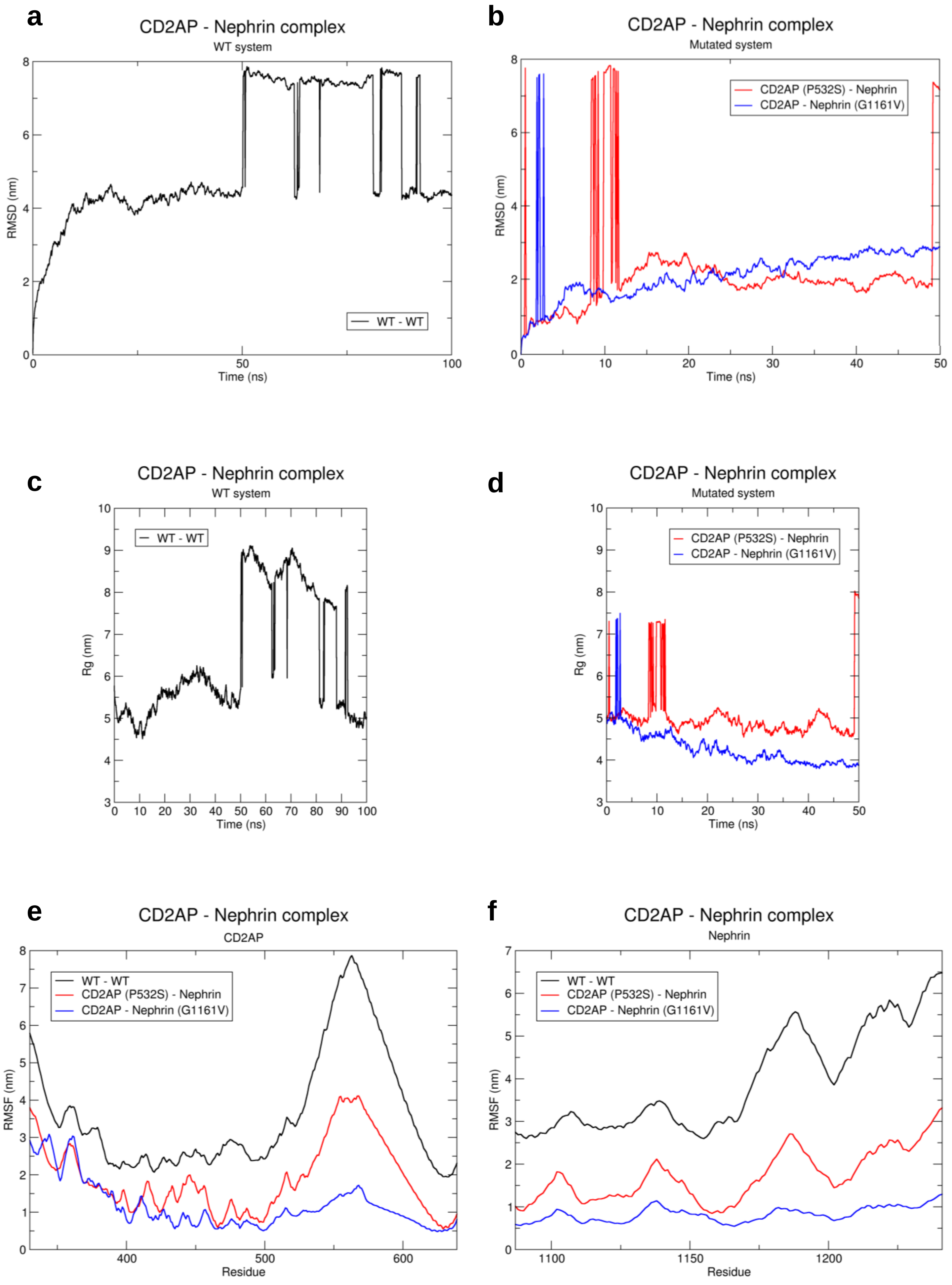

### Supplementary figure 4

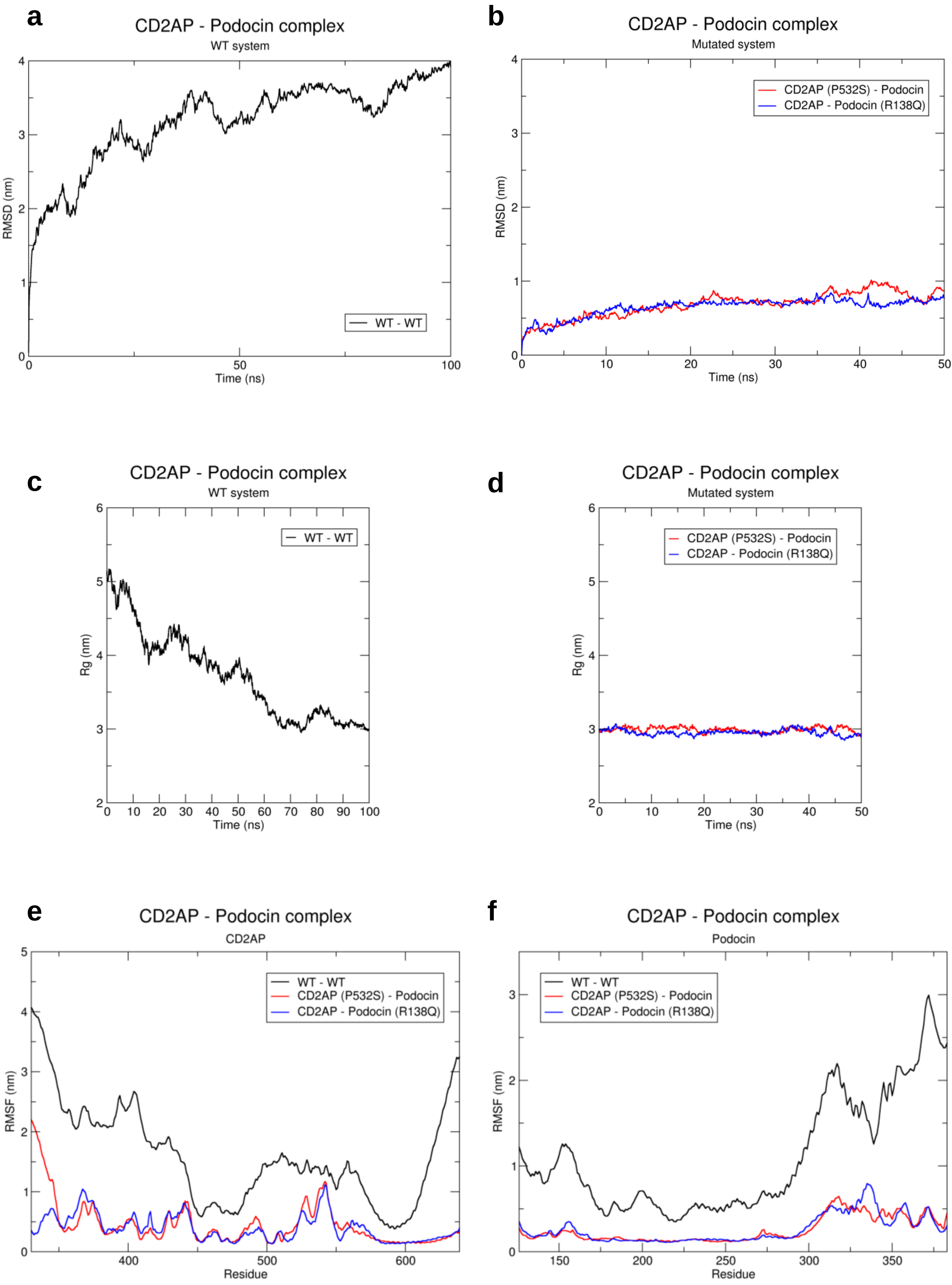

### Supplementary figure 5

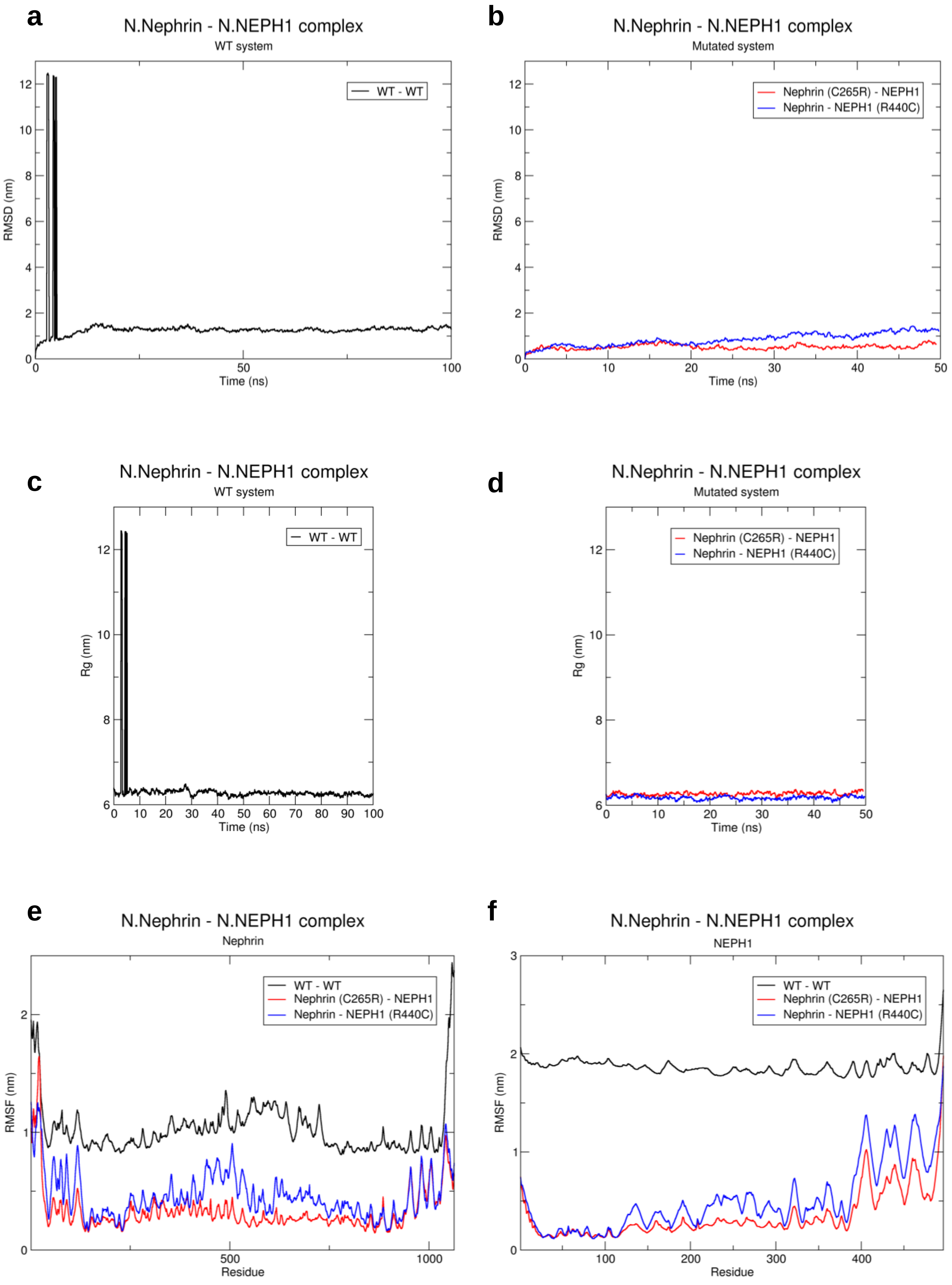

### Supplementary figure 6

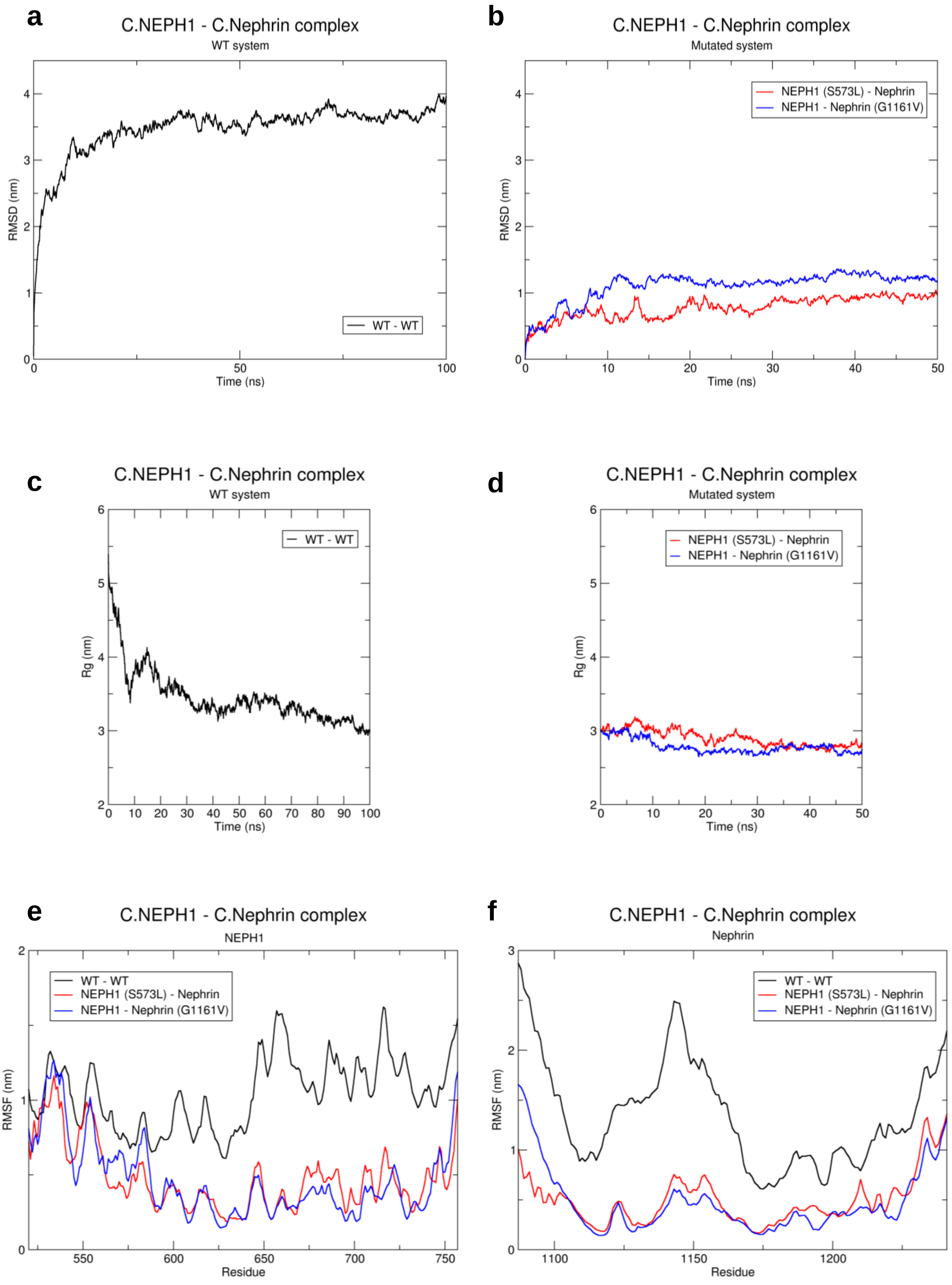

### Supplementary figure 7

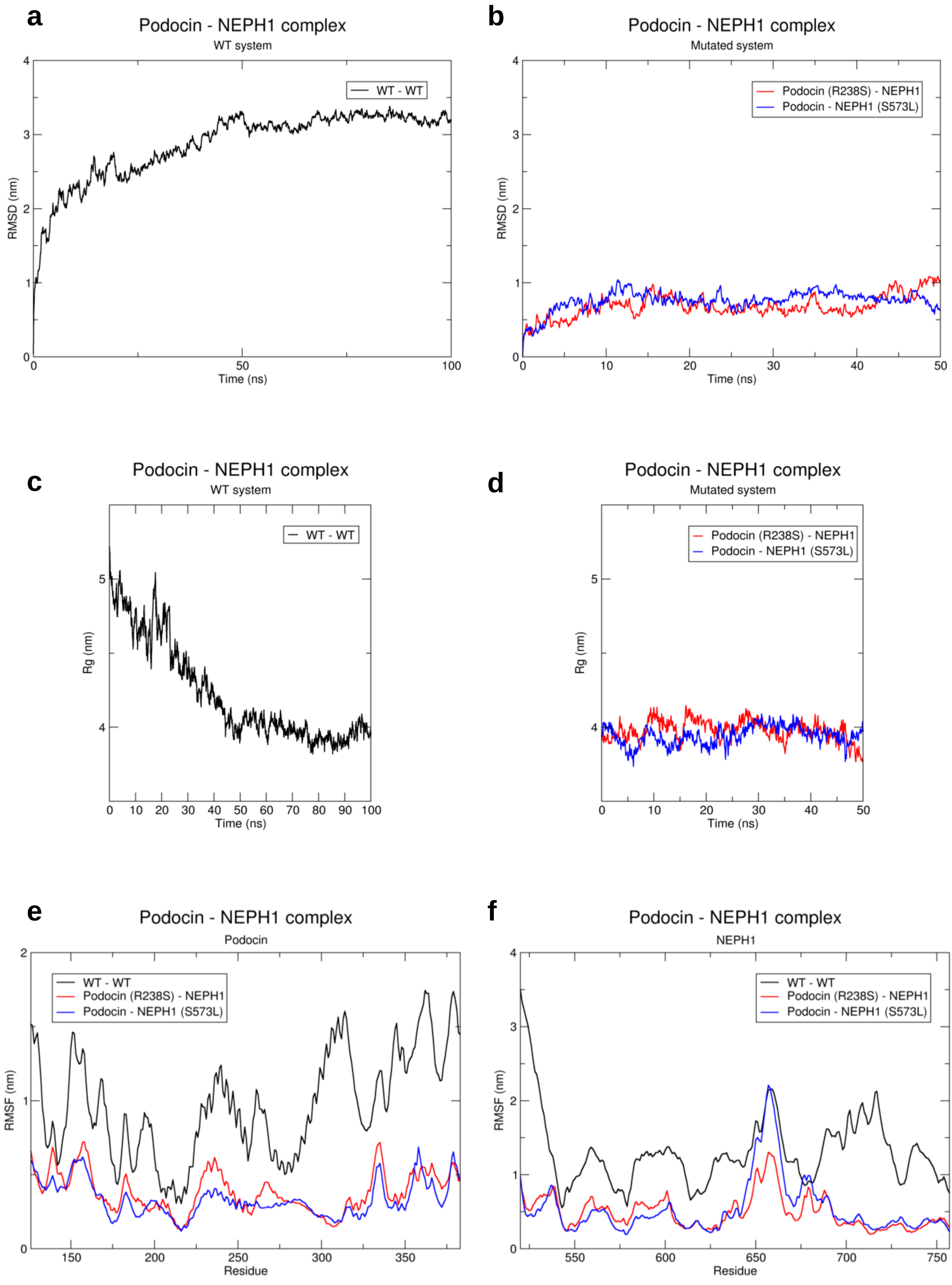

### Supplementary figure 8

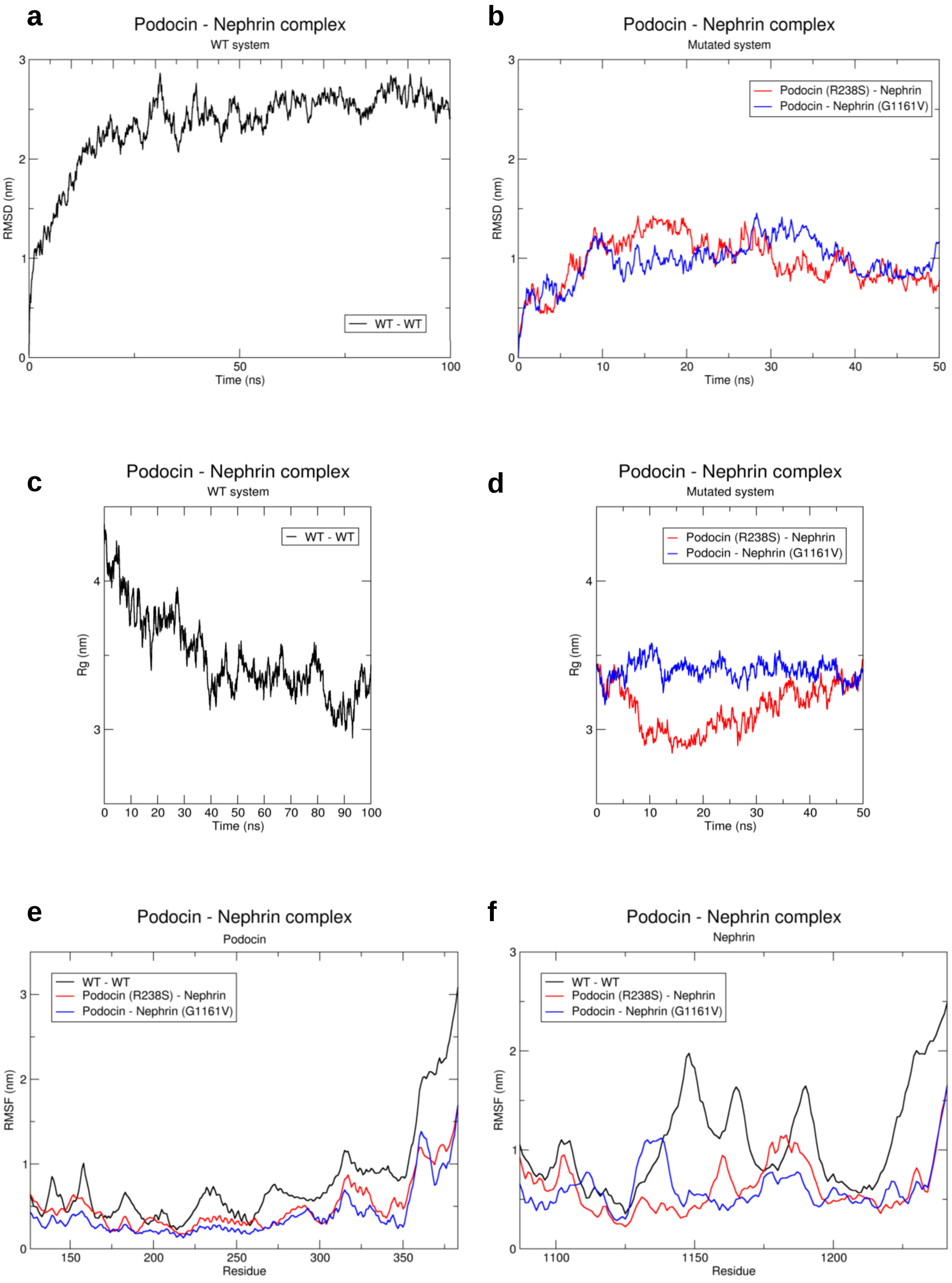

### Supplementary figure 9

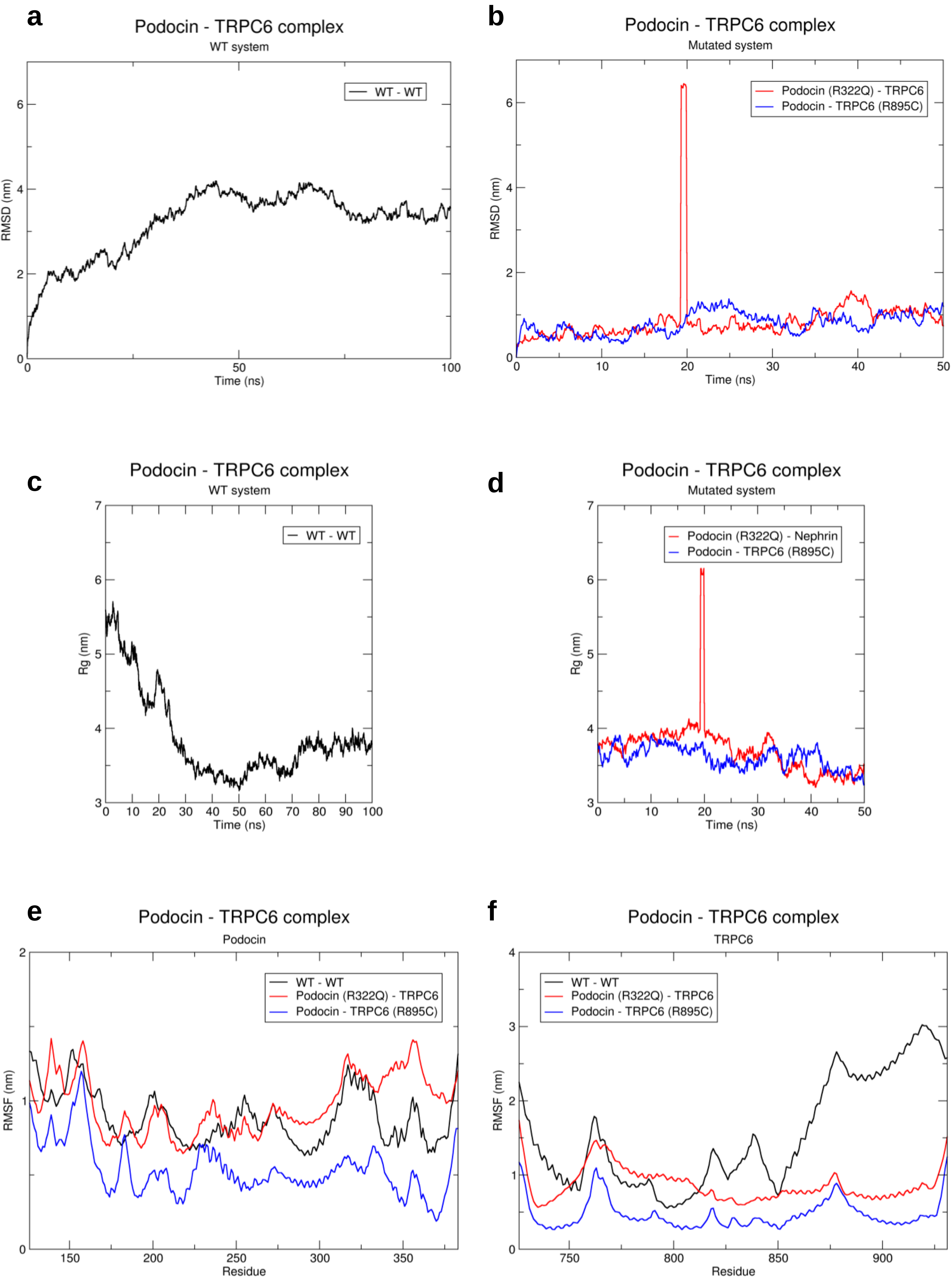

**Supplementary figure 10**

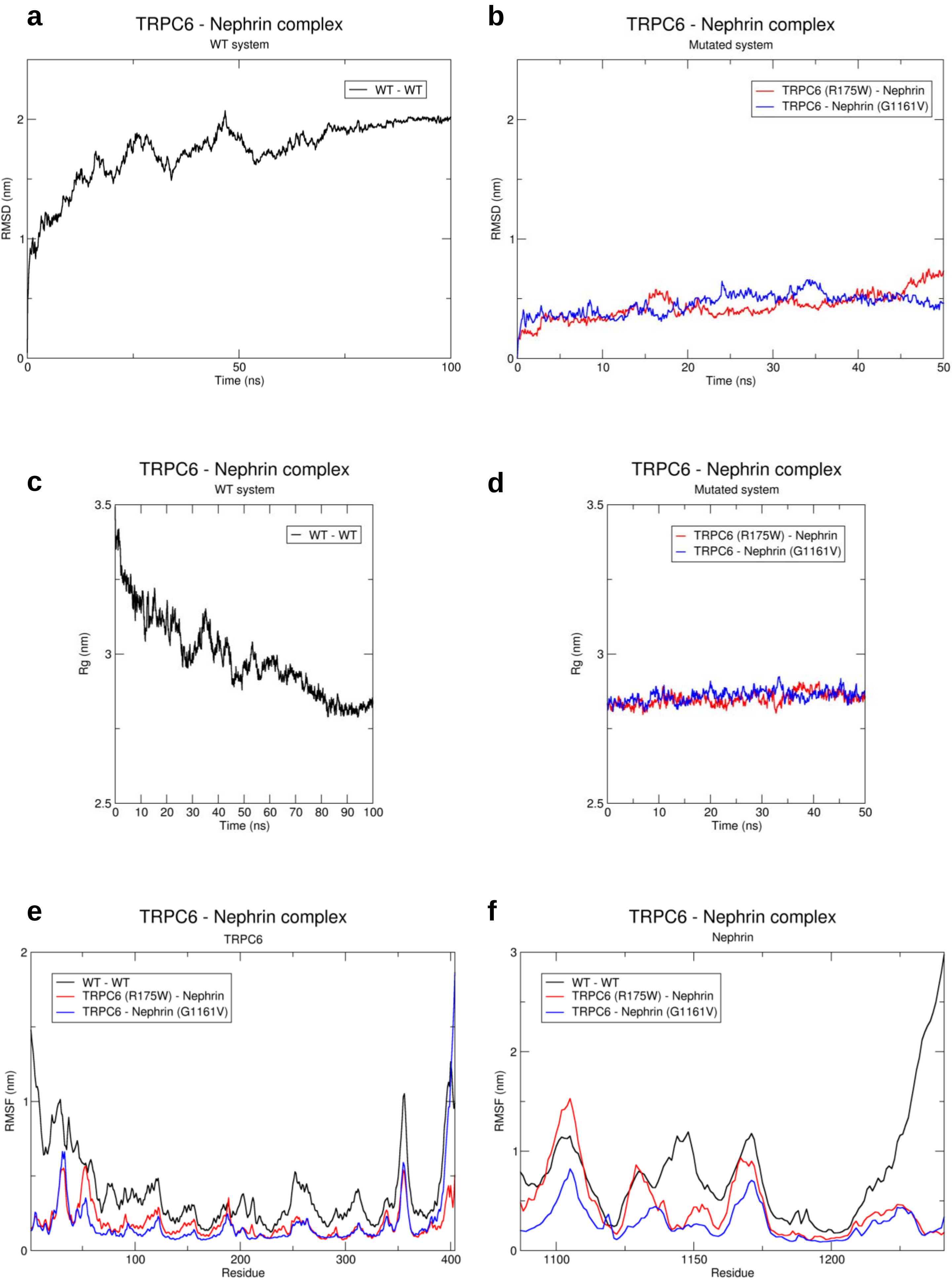

#### Supplementary Figure Legends

##### Supplementary Figure 1: Quality assessment of AlphaFold-predicted models

**a.** Podocin: The predicted aligned error (PAE) plot indicates high accuracy in the PHB domain, while deviations are observed in its relative positioning to the disordered C-terminal region. **b.** The PAE plot demonstrates high confidence in predicting the structured regions (three SH3 domains and the coiled-coil domain), although their relative positioning is uncertain due to disordered linking regions. **c.** NEPH1: The five Ig domains are accurately predicted and maintain reliable relative positions. However, the transmembrane region, though well-predicted, shows poor positional alignment with the Ig domains. Uncertainty increases in the cytoplasmic domain. **d.** TRPC6: The model is generally accurate, except for disordered stretches in the N-terminal and C-terminal regions. **e.** Nephrin: The Ig and transmembrane domains are accurately predicted with good positional consistency, though the disordered cytoplasmic domain lacks conformational stability. **f.** HSP27: The ordered HSP domain is predicted with high confidence and is well-positioned relative to the disordered N- and C-terminal regions exhibiting significant deviations. Overall, the analysis shows high confidence (pLDDT scores >85%) for structured domains across all target proteins, while low confidence (<55%) is noted for the intrinsically disordered regions (IDRs).

##### Supplementary Figure 2: Average energy plots of all protein-protein interactions

The end-state free energy of bindings of all the protein-protein complexes in WT (averaged over 100 ns) and mutated (averaged over 50 ns) states were calculated using molecular mechanics with the generalized Born and surface area solvation (MM-GBSA) method. **a.** CD2AP-nephrin complex. **b.** CD2AP-podocin complex. **c.** Extracellular nephrin-NEPH1 complex. **d.** Cytoplasmic NEPH1-nephrin complex. **e.** Podocin-NEPH1 complex. **f.** Podocin-nephrin complex. **g.** Podocin-TRPC6 complex. **h.** TRPC6-nephrin complex.

**Supplementary Figure 3: Details of the WT and mutated CD2AP-nephrin trajectories**

**a.** RMSD plot of the WT CD2AP-nephrin trajectory (100 ns). **b.** RMSD plot of the mutated CD2AP-nephrin trajectories (50 ns each). **c.** Rg plot of the WT CD2AP-nephrin trajectory (100 ns). **d.** Rg plot of the mutated CD2AP-nephrin trajectories (50 ns each). RMSF plots of **e.** CD2AP and **f.** nephrin in the WT and mutated trajectories.

**Supplementary Figure 4: Details of the WT and mutated CD2AP-podocin trajectories**

**a.** RMSD plot of the WT CD2AP-podocin trajectory (100 ns). **b.** RMSD plot of the mutated CD2AP-podocin trajectories (50 ns each). **c.** Rg plot of the WT CD2AP-podocin trajectory (100 ns). **d.** Rg plot of the mutated CD2AP-podocin trajectories (50 ns each). RMSF plots of **e.** CD2AP and **f.** podocin in the WT and mutated trajectories.

**Supplementary Figure 5: Details of the WT and mutated extracellular nephrin-NEPH1 trajectories**

**a.** RMSD plot of the WT extracellular nephrin-NEPH1 trajectory (100 ns). **b.** RMSD plot of the mutated extracellular nephrin-NEPH1 trajectories (50 ns each). **c.** Rg plot of the WT extracellular nephrin-NEPH1 trajectory (100 ns). **d.** Rg plot of the mutated extracellular nephrin-NEPH1 trajectories (50 ns each). RMSF plots of extracellular **e.** nephrin and **f.** NEPH1 in the WT and mutated trajectories.

**Supplementary Figure 6: Details of the WT and mutated cytoplasmic NEPH1-nephrin trajectories**

**a.** RMSD plot of the WT cytoplasmic NEPH1-nephrin trajectory (100 ns). **b.** RMSD plot of the mutated cytoplasmic NEPH1-nephrin trajectories (50 ns each). **c.** Rg plot of the WT cytoplasmic NEPH1-nephrin trajectory (100 ns). **d.** Rg plot of the mutated cytoplasmic

NEPH1-nephrin trajectories (50 ns each). RMSF plots of cytoplasmic **e.** NEPH1 and **f.** nephrin in the WT and mutated trajectories.

**Supplementary Figure 7: Details of the WT and mutated podocin-NEPH1 trajectories**

**a.** RMSD plot of the WT podocin-NEPH1 trajectory (100 ns). **b.** RMSD plot of the mutated podocin-NEPH1 trajectories (50 ns each). **c.** Rg plot of the WT podocin-NEPH1 trajectory (100 ns). **d.** Rg plot of the mutated podocin-NEPH1 trajectories (50 ns each). RMSF plots of **e.** podocin and **f.** NEPH1 in the WT and mutated trajectories.

**Supplementary Figure 8: Details of the WT and mutated podocin-nephrin trajectories**

**a.** RMSD plot of the WT podocin-nephrin trajectory (100 ns). **b.** RMSD plot of the mutated podocin-nephrin trajectories (50 ns each). **c.** Rg plot of the WT podocin-nephrin trajectory (100 ns). **d.** Rg plot of the mutated podocin-nephrin trajectories (50 ns each). RMSF plots of **e.** podocin and **f.** nephrin in the WT and mutated trajectories.

**Supplementary Figure 9: Details of the WT and mutated podocin-TRPC6 trajectories**

**a.** RMSD plot of the WT podocin-TRPC6 trajectory (100 ns). **b.** RMSD plot of the mutated podocin-TRPC6 trajectories (50 ns each). **c.** Rg plot of the WT podocin-TRPC6 trajectory (100 ns). **d.** Rg plot of the mutated podocin-TRPC6 trajectories (50 ns each). RMSF plots of **e.** podocin and **f.** TRPC6 in the WT and mutated trajectories.

**Supplementary Figure 10: Details of the WT and mutated TRPC6-nephrin trajectories**

**a.** RMSD plot of the WT TRPC6-nephrin trajectory (100 ns). **b.** RMSD plot of the mutated TRPC6-nephrin trajectories (50 ns each). **c.** Rg plot of the WT TRPC6-nephrin trajectory (100 ns). **d.** Rg plot of the mutated TRPC6-nephrin trajectories (50 ns each). RMSF plots of **e.** TRPC6 and **f.** nephrin in the WT and mutated trajectories.

**Movie legends**

**Movie S1: MD trajectory of CD2AP (magenta) – nephrin (blue) complex.** The simulation time of the WT trajectory was 100 ns. The WT trajectory revealed that the proteins interacted directly and transiently. The trajectories of modified systems, initiated from the endpoint of the WT trajectory, were simulated for 50 ns, with a mutation in each protein. Nephrin's mutation promoted self-disorder at the N′ terminal by altering the secondary structure from helix to coil. The depiction showcases the proteins in cartoon form, highlighting the mutated residue in licorice representation (*P532S* – green and *G1161V* – green). The VMD-rendered movie excluded water and ions for enhanced visualization. In this complex, we used truncated CD2AP (330 – 639) and the cytoplasmic region of nephrin (1087 – 1241).

**Movie S2: MD trajectory of CD2AP (magenta) – podocin (red) complex.** The simulation time of the WT trajectory was 100 ns. We observed substantial folding of CD2AP's coiled-coil domain in contact with podocin. The trajectories of modified systems, initiated from the endpoint of the WT trajectory, were simulated for 50 ns, with a mutation in each protein. We observed stable interactions with minimal changes in the native interacting pattern. The depiction showcases the proteins in cartoon form, highlighting the mutated residue in licorice representation (*P532S* – green and *R138Q* – green). The VMD-rendered movie excluded water and ions for enhanced visualization. In this complex, we used truncated CD2AP (330 – 639) and podocin (126 – 383).

**Movie S3: MD trajectory of N.Nephrin (blue) – N.NEPH1 (green) complex.** The simulation time of the WT trajectory was 100 ns. The extracellular regions of nephrin and NEPH1 from adjacent foot processes exhibited stable complexation via Ig domains. The trajectories of modified systems, initiated from the endpoint of the WT trajectory, were simulated for 50 ns, with a mutation in each protein. The modified systems exhibited stable complexation. The depiction showcases the proteins in cartoon form, highlighting the mutated residue in licorice representation (*C265R* – red and *R440C* – red). The VMD-rendered movie excluded water and ions for enhanced visualization. In this complex, we used the extracellular regions of nephrin (1 – 1063) and NEPH1 (1 – 496).

**Movie S4: MD trajectory of C.NEPH1 (green) – C.Nephrin (blue) complex.** The simulation time of the WT trajectory was 100 ns. The cytoplasmic regions of NEPH1 and nephrin interacted via IDRs. The trajectories of modified systems, initiated from the endpoint of the WT trajectory, were simulated for 50 ns, with a mutation in each protein. Both mutations enable the complex to form more interactions. The depiction showcases the proteins in cartoon form, highlighting the mutated residue in licorice representation (*S573L* – red and *G1161V* – red). The VMD-rendered movie excluded water and ions for enhanced visualization. In this complex, we used the cytoplasmic regions of NEPH1 (520 – 757) and nephrin (1087 – 1241).

**Movie S5: MD trajectory of podocin (red) – NEPH1 (green) complex.** The simulation time of the WT trajectory was 100 ns. NEPH1 altered the conformation of podocin slightly by influencing the IDRs present in podocin's interacting interface. The trajectories of modified systems, initiated from the endpoint of the WT trajectory, were simulated for 50 ns, with a mutation in each protein. Mutations in each protein (podocin – *R238S* and NEPH1 – *S573L*) stabilized the complex, enhancing NEPH1's interacting interface. The depiction showcases the proteins in cartoon form, highlighting the mutated residue in licorice representation (*R238S* – blue and *S573L* – blue). The VMD-rendered movie excluded water and ions for enhanced visualization. In this complex, we used truncated podocin (126 – 383) and the cytoplasmic region of NEPH1 (520 – 757).

**Movie S6: MD trajectory of podocin (red) – nephrin (blue) complex.** The simulation time of the WT trajectory was 100 ns. Nephrin influenced the IDRs within podocin's interacting interface and facilitated the folding of its ordered regions by inducing a distinct bend towards the end of the PHB domain. The trajectories of modified systems, initiated from the endpoint of the WT trajectory, were simulated for 50 ns, with a mutation in each protein. Mutation in each protein had minimal effect on the sizes of the interacting interfaces, but we observed an increase in the number of interactions. The depiction showcases the proteins in cartoon form, highlighting the mutated residue in licorice representation (*R238S* – green and *G1161V* – green). The VMD-rendered movie excluded water and ions for enhanced visualization. In this complex, we used truncated podocin (126 – 383) and the cytoplasmic region of nephrin (1087 – 1241).

**Movie S7: MD trajectory of podocin (red) – TRPC6 (purple) complex.** The simulation time of the WT trajectory was 100 ns. IDRs in the interacting interface of podocin were instrumental in facilitating the movement and conformational changes in TRPC6. The trajectories of modified systems, initiated from the endpoint of the WT trajectory, were simulated for 50 ns, with a mutation in each protein. *R322Q* resulted in improved interactions in the complex. The depiction showcases the proteins in cartoon form, highlighting the mutated residue in licorice representation (*R322Q* – green and *R895C* – green). The VMD-rendered movie excluded water and ions for enhanced visualization. In this complex, we used truncated podocin (126 – 383) and the C-terminal region of TRPC6 (726 – 931).

**Movie S8: MD trajectory of TRPC6 (purple) – nephrin (blue) complex.** The simulation time of the WT trajectory was 100 ns. The extended conformation of nephrin became compact while TRPC6 remained stable. The trajectories of modified systems, initiated from the endpoint of the WT trajectory, were simulated for 50 ns, with a mutation in each protein. Mutation in each protein maintained the interacting regions of both proteins, which were almost identical to the native interacting regions while improving the number of interactions. The depiction showcases the proteins in cartoon form, highlighting the mutated residue in licorice representation (*R175W* – red and *G1161V* – red). The VMD-rendered movie excluded water and ions for enhanced visualization. In this complex, we used the N-terminal region of TRPC6 (1 – 404) and the cytoplasmic region of nephrin (1087 – 1241).
